# Mapping allosteric rewiring in related protein structures from collections of crystallographic multiconformer models

**DOI:** 10.1101/2025.05.23.655529

**Authors:** Akshay Raju, Shivani Sharma, Blake T. Riley, Shakhriyor Djuraev, Yingxian Tan, Minyoung Kim, Toufique Mahmud, Daniel A. Keedy

## Abstract

How do related proteins with a common fold perform diverse biological functions? Although the average structure may be similar, structural excursions from this average may differ, giving rise to allosteric rewiring that enables differential activity and regulation. However, this idea has been difficult to test in detail. Here we used the qFit algorithm to model “hidden” alternate conformations from electron density maps for an entire protein family, the Protein Tyrosine Phosphatases (PTPs), spanning 26 enzymes and 221 structures. To interrogate these multiconformer models, we developed a new algorithm, Residue Interaction Networks From Alternate conformations In RElated structures (RINFAIRE), that calculates networks of interactions between flexible residues and quantitatively compares them. We show that PTPs share a common allosteric network which rewires dynamically in response to catalytic loop motions or active-site vs. allosteric ligand binding, but also that individual PTPs have unique allosteric signatures. As experimental validation, we show that targeted mutations at residues with varying sequence conservation but high network connectivity modulate enzyme catalysis, including a surprising enhancement of activity. Overall, our work provides new tools for understanding how evolution has recycled modular macromolecular building blocks to diversify biological function. RINFAIRE is available at https://github.com/keedylab/rinfaire.

## Introduction

Allostery is a prevalent regulatory mechanism in biology ^1,2^, allowing proteins to respond to stimuli such as ligand binding at one site by altering their structure and function at another site. Allosteric communication within a protein fold ^3,4^ may occur through a variety of mechanisms ^5^, including conformational rearrangements of loops and linkers ^6,7^, shifting networks of side-chain interactions ^8^, and changes in dynamics with an unchanged average conformation ^9^.

A key question in molecular biophysics is whether allosteric wiring either (i) is a property of a protein fold and thus conserved over evolution or (ii) differs between related proteins to diversify regulation/function. Some lines of evidence point to conservation of allostery: statistical coupling analysis (SCA) of coevolving amino acids reveals sectors of residues ^10^ that highlight allosteric sites ^11^, and conformational dynamics are somewhat conserved within a protein fold even when sequence diverges ^12^. However, smaller-scale fast dynamics, which may play roles in diversifying function, are more divergent than larger-scale slow dynamics ^13^. Altogether, it remains unclear to what extent functionally relevant allosteric wiring is customized in different homologs within the constraints of a common fold.

This question is relevant to many protein families, not least of which are the protein tyrosine phosphatase (PTP) enzymes. Many PTPs are validated therapeutic targets for diabetes, obesity, cancer, autoimmune diseases, and neurological diseases ^14^. Here we focus on the ∼37 ^15^ class I classical PTPs which are specific for phosphotyrosine (pTyr) moieties in substrate proteins ^15,16^. Despite structural conservation of the PTP catalytic domain (**Fig. 1**), the average sequence identity is only 34.4% (range: 21.7–98.5%), indicating substantial divergence that may manifest as rewired allostery. Consistent with this idea, distinct regulatory domains in different PTPs ^17^ have been shown to regulate the catalytic domain in unique ways, including the α7 helix in PTP1B (PTPN1) ^18–21^ and in the closely related TCPTP (PTPN2) ^22^, the N-terminal autoinhibitory SH2 domains in SHP2 (PTPN11) ^23–25^, and the non-catalytic PTP-like D2 domain in certain receptor-type PTPs ^26,27^. Crystal structures of different PTPs also reveal distinct patterns of surface features ^16,26^, suggesting the existence of unique, non-orthosteric binding sites. Indeed, early small-molecule allosteric modulators ^28^ have been reported for PTP1B ^18,21,29–31^, SHP2 ^32,33^, and STEP (PTPN5) ^34^.

**Figure 1:**
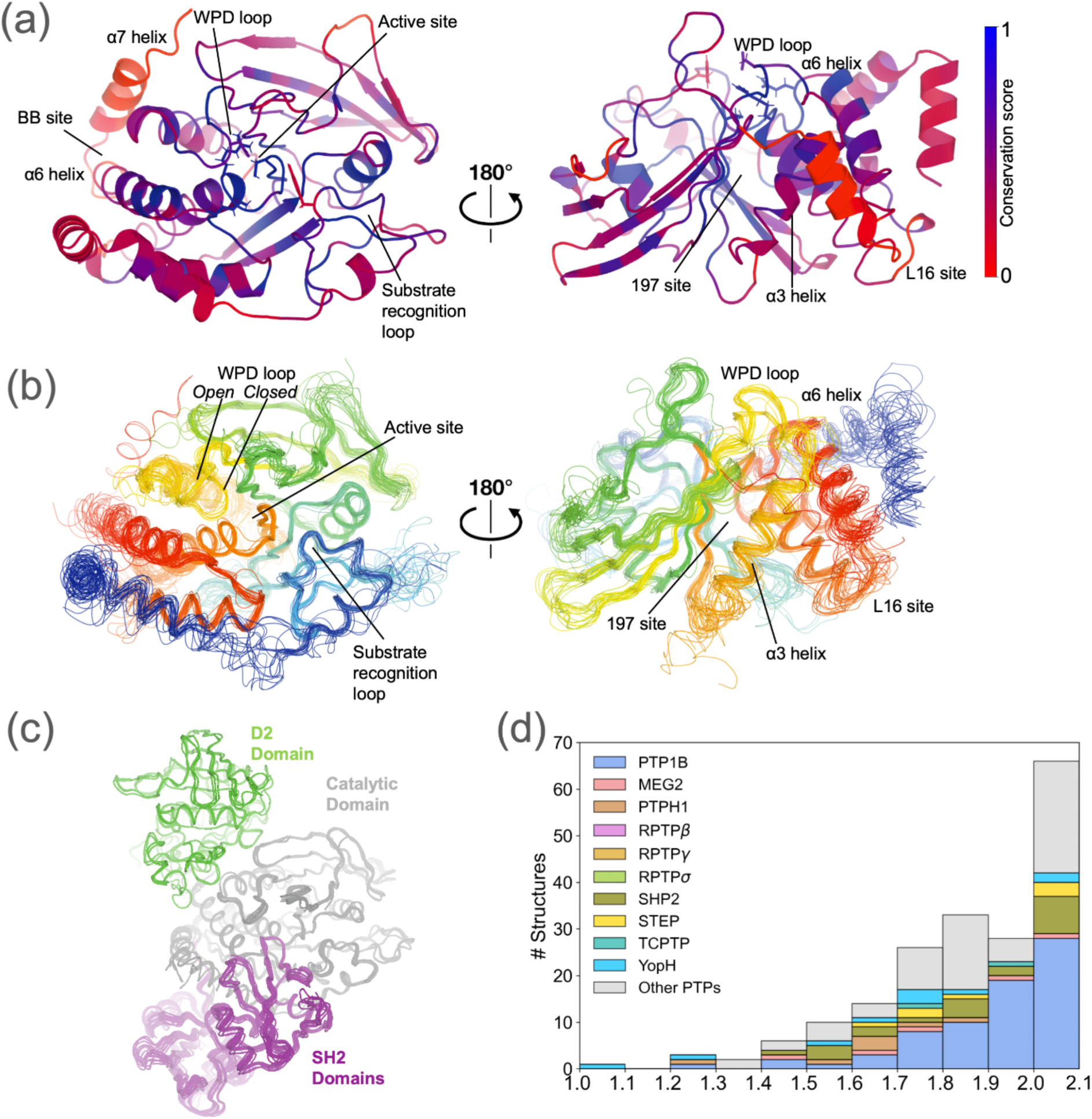
Overview of crystal structures of related PTP enzymes. This study uses a dataset of 170 publicly available PTP crystal structures with sufficiently high resolution (≤ 2.1 Å), representing 26 distinct PTP enzymes. **(a)** Sequence conservation from a structure-based sequence alignment (see Methods), mapped to a representative structure of the catalytic domain of the archetypal PTP family member, PTP1B (PDB ID: 1sug)^54^. Key sites are indicated, such as active-site loops and allosteric sites; catalytic residues (Asp181, Arg221, Cys215, Gln262, Tyr46; PTP1B numbering) are shown as sticks. **(b)** Structural alignment using Cα backbone atoms for all PTP structures studied here, colored from N-terminus (blue) to C-terminus (red). **(c)** Structural alignment using the shared catalytic domain (gray) for all PTP structures studied here that contain additional ordered protein domains: C-terminal non-catalytic “D2” domains (green), or N-terminal SH2 domains (purple). **(d)** Resolution distribution for all available PTP structures (with resolution ≤ 2.1 Å cutoff). Different colors indicate different PTP enzymes. Each bin is left-inclusive and right-exclusive except the last bin with both inclusive (structures at 2.1 Å are included).

Despite this promising outlook, many mysteries remain about the evolutionary divergence of allosteric wiring in the PTP catalytic domain. For example, SCA suggested two allosteric sectors shared among PTPs ^35^, but MD (molecular dynamics) analysis suggested divergent allostery based on differences in correlated structural motions ^36^. Increased clarity about allosteric similarities vs. differences among PTPs would aid in developing allosteric modulators that are specific for individual PTPs, helping these enzymes shed their reputation of being “undruggable” ^37^.

Previously, several approaches have been used to elucidate allosteric wiring in related proteins like PTPs. SCA ^10,11^ generates testable hypotheses about allosteric sectors, but questions remain about the physical interpretation of these sectors. Computational structure-based methods to study allostery ^5^ including MD simulations ^36,38–41^, normal mode analysis (NMA) ^42^, and machine learning ^43^ are often too computationally intensive to scale well to large protein families and/or rely on simplified force fields. Experiments like nuclear magnetic resonance (NMR) spectroscopy, cryo-electron microscopy (cryo-EM), and site-directed mutagenesis provide direct insights into dynamics and function ^20,44^, but have limited throughput and spatial resolution.

Bridging computation and experimental data, multiconformer modeling from crystallographic electron density maps with qFit (**Fig. 2a**) ^45–49^ yields parsimonious alternate conformations of protein side-chain and backbone atoms. qFit models are consistent with NMR dynamics data ^50^ and reveal entropic compensation mechanisms from ligand binding ^51^. To analyze the complex coupling between spatially adjacent alternate conformations in qFit models, previously the CONTACT algorithm used steric clashes only; the resulting networks were validated by NMR for the model enzyme dihydrofolate reductase (DHFR) ^8^ and revealed a ligand-dependent signaling mechanism for mPGES-1 ^52^. However, CONTACT does not consider interaction types beyond steric clashes, nor does it offer machinery to compare networks for related structures, leaving key gaps in its capabilities.

**Figure 2:**
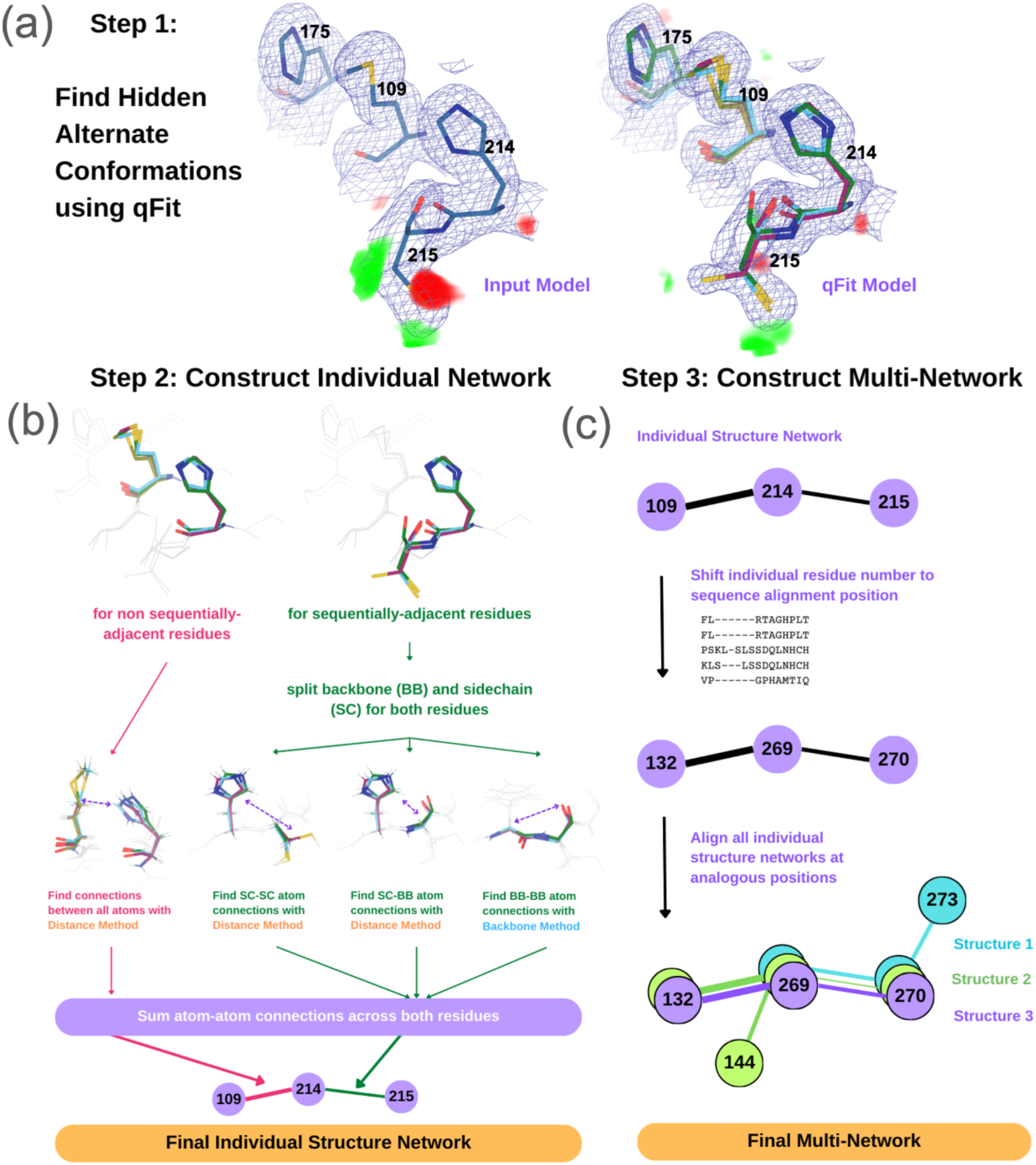
RINFAIRE workflow to generate multinetworks from related qFit multiconformer models. **(a)** qFit multiconformer modeling for each structure identifies “hidden” alternate conformations that better explain the electron density. In this example (PDB ID: 3eax) ^57^, qFit finds previously unmodeled alternate conformations for the catalytic Cys215 and several spatially adjacent residues near the active site of PTP1B. 2Fo-Fc density contoured at 1 σ (blue mesh); Fo-Fc difference density contoured at +/- 3 σ (green/red volumes). **(b)** The qFit multiconformer model is used by RINFAIRE to construct an individual structure network. This example features interactions between selected residues from panel (a). See also **Fig. S2.** **(c)** All of the individual networks are aligned using a structure-based sequence alignment, generating a “multinetwork”. In subsequent steps, an overall sum network can be computed, or sum networks composed of subsets of structures can be compared.

To fill these gaps, we have developed a new algorithm, **R**esidue **I**nteraction **N**etworks **F**rom **A**lternate conformations In **RE**lated structures (RINFAIRE). By using a distance-based approach, RINFAIRE implicitly captures a wider range of interactions between alternate conformations, including unfavorable steric clashes as well as favorable hydrogen bonds (H-bonds), van der Waals packing, and ionic interactions. RINFAIRE also aligns and scales residue interaction networks (RINs) from multiple input qFit models, subsets these RINs based on custom metadata, and quantitatively compares different sum networks corresponding to distinct subsets of structures. We have deployed our novel qFit + RINFAIRE computational pipeline to study allosteric networks for all structurally characterized PTPs. Leveraging the growth of the Protein Data Bank (PDB) ^53^, we studied 221 PTP catalytic domain structures spanning 26 distinct PTP enzymes. Our results reveal how allosteric wiring in the PTP catalytic domain changes between well-known global conformational states relevant to catalysis, upon binding to active-site vs. allosteric ligands relative to the apo state, and in different PTPs with distinct functional and/or regulatory properties. RINFAIRE is free and open-source software, available at https://github.com/keedylab/rinfaire.

## Results

### Creating a dataset of multiconformer models of the PTP family

After filtering by resolution (≤ 2.1 Å) and automated re-refinement (see Methods), we assembled 170 high-resolution crystal structures of PTPs, representing 26 distinct human PTPs plus another 6 orthologous PTPs from other species (**Fig. S1**). PTP1B (PTPN1) is the most represented PTP, followed by SHP2 (PTPN11) and bacterial YopH (**Fig. 1d**).

The PTP structures in our dataset have a substantial degree of sequence and structural conservation across the catalytic domain, especially near the active site (**Fig. 1a**). The average sequence conservation value for residue positions that align with the PTP1B catalytic domain is 49.8%. Important loops for catalysis such as the P loop, WPD loop, Q loop, and substrate recognition loop (i.e. pTyr binding loop) have especially high conservation across our structural dataset, having average conservation of 90.7%, 75.8%, 70.4%, and 70.0% respectively.

Many of the regions that are well conserved in terms of sequence are also well conserved in terms of structure (**Fig. 1b**). The backbone in the catalytic domain shows relatively little variation overall and for the active-site P loop, Q loop, and substrate recognition loop (**Fig. 1b**). A notable exception is the dynamic WPD loop ^21,55^ which clusters into three distinct states: predominantly the canonical open conformation and closed conformation, with a few examples of an atypically open or super-open conformation in a few PTPs such as STEP (PTPN5) and YopH. Additional domains exist that are unique to some PTPs, including SH2 in SHP1/SHP2 and the inactive catalytic domain D2 in some receptor-type PTPs (**Fig. 1c**) ^17,26,56^.

### Using alternate conformations to generate residue interaction networks

Protein crystallographic electron density maps often reveal “hidden” alternate conformations that are unmodeled in the publicly available structures ^58^. To better represent the structural heterogeneity present in our dataset, we used the automated multiconformer modeling algorithm qFit ^45–49^ for all 170 crystal structures in our dataset (**Fig. 2a**). qFit increased the average number of alternate conformations by 17.7% (from 1.0 to 1.2 conformations per residue). Based on R_free_ and R-gap (R_free_-R_work_) (**Fig. S3**), qFit adds alternate conformations that help explain the experimental data better than the original deposited structures and do not overfit the data.

Of the 170 structures, 50 have non-crystallographic symmetry with multiple non-identical instances of the PTP catalytic domain. Following qFit refinement, we separated these instances, resulting in 221 distinct catalytic domain structures for subsequent analysis.

The qFit models contain many instances of coupled alternate conformations at important sites in the structurally conserved catalytic domain. For example, a deposited structure of the archetypal family member PTP1B (PDB ID: 3eax) had a missing alternate conformation at the catalytic cysteine (Cys215), as indicated by difference electron density (**Fig. 2a**). qFit successfully modeled this new rotamer conformation, along with subtle alternate conformations of a sequentially neighboring residue and several spatially adjacent residues (His214, Met109, His175) in a β sheet, resulting in diminished difference density (**Fig. 2a**).

We next sought to compute the network of such interactions in each qFit model and compare them across all our PTP structures. To do so, we developed a new computational method called **R**esidue **I**nteraction **N**etworks **F**rom **A**lternate conformations **I**n **RE**lated structures, or RINFAIRE (**Fig. 2b-c**). RINFAIRE proceeds in two main stages. First, in the individual-network stage, a RIN is generated for each structure based on interactions between alternate-conformation atoms in residues that are either adjacent in space or adjacent in sequence (**Fig. 2b**). In contrast to past methods for computing RINs from multiconformer models that only modeled repulsive steric clashes ^8^, RINFAIRE implicitly incorporates favorable van der Waals forces, hydrogen bonds, ionic bonds, and other local interactions (albeit in a coarse-grained fashion). Second, in the multinetwork stage, individual RINs are aligned based on a structure-based multiple sequence alignment, allowing analogous residues to be directly compared across all networks (**Fig. 2c**). Once aligned, the networks are log normalized to account for differences in numbers of alternate conformations (e.g. due to resolution differences) and prepared for comparative analyses including summation per edge and calculation of differences between defined subsets (see Methods).

### The consensus allosteric network of the PTP catalytic domain

To map allosteric connections that are most represented across PTPs, we used RINFAIRE to generate a sum network for all PTP structures in our dataset. We identified the most structurally conserved components of this network by restricting to the top 5% of edges (by edge weight), resulting in a pruned sum network of 89 nodes and 120 edges with a cyclical topology (**Fig. 3a-b**). To identify important residues in this network, we used weighted degree, i.e. the sum of edge weights for each node. Hereafter in this manuscript, we refer to weighted degree as simply degree.

**Figure 3:**
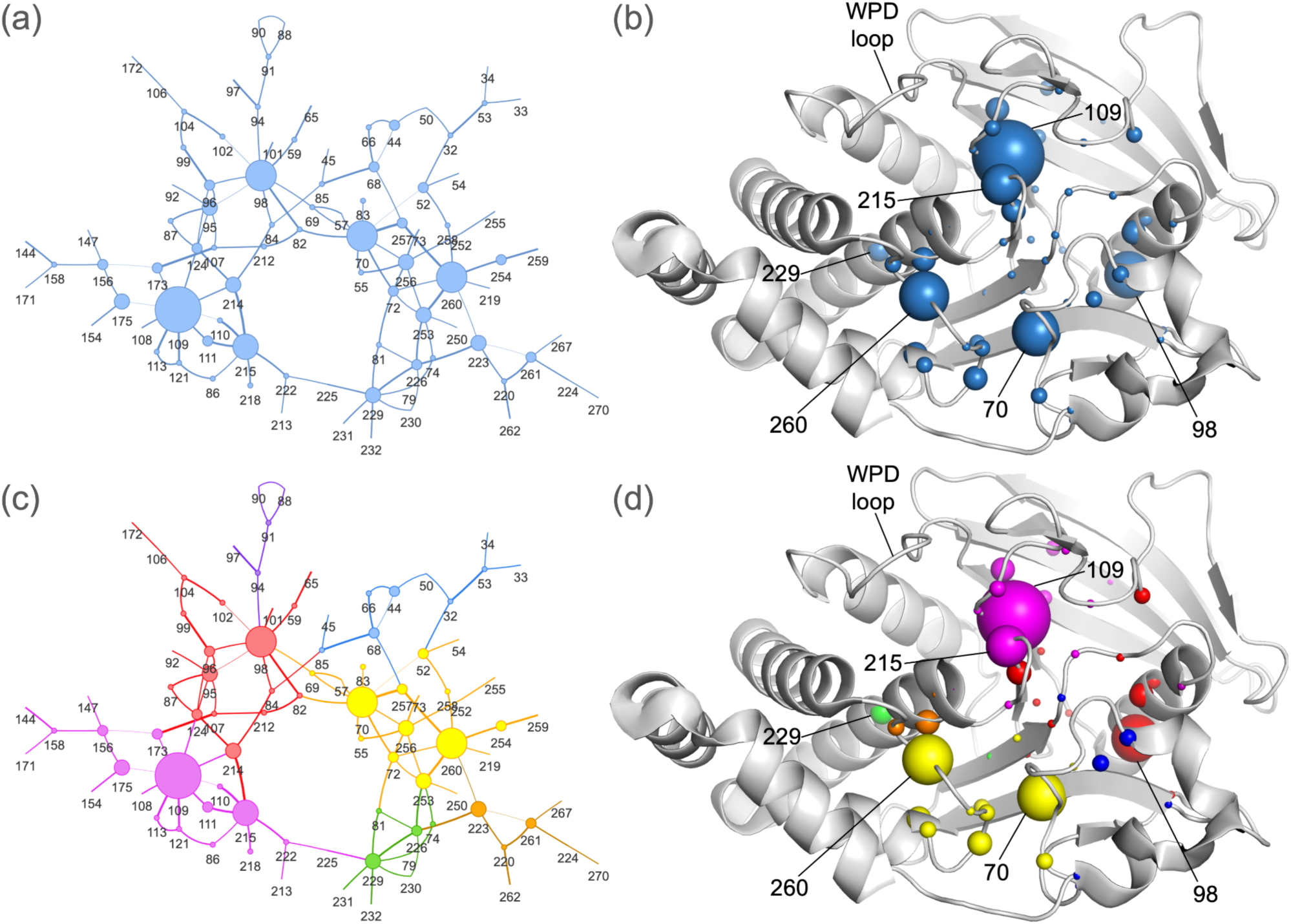
Sum network analysis using all PTP structures. **(a)** 2D diagram of the RINFAIRE sum network for all suitable PTP structures, showing the top 5% of edges based on edge weight. Line thickness represents edge weight; node size represents degree. Sets of nodes with less than 5 edges are hidden for visual clarity. **(b)** Sum network mapped onto a structure of PTP1B (PDB ID: 1t49) ^18^. Sphere size represents degree. The archetypal family member PTP1B is used as a reference for residue labeling; only those residues with an analogous residue in PTP1B are shown. **(c)** The sum network is partitioned into 7 distinct communities (colors) using the Girvan-Newman algorithm (see Methods and **Fig. S4**). **(d)** The communities are mapped onto a representative structure of PTP1B (PDB ID: 1t49).

Met109 (PTP1B numbering) has the highest degree overall. Although this residue has not been previously highlighted as key to catalysis, it is 100% conserved across human PTP sequences ^17^, and in our network is connected to several residues that bridge to the catalytic Cys215 (**Fig. 2a**), the active-site E loop, and the N-terminal hinge point of the catalytic WPD loop (**Fig. 2a**), whose dynamic motions are critical for catalysis in PTPs ^59^. The next highest-degree residues are at key functional sites and/or exhibit dynamic behavior (**Fig. 3a-b**). Ser70 is near the substrate recognition loop and P loop, in a dynamic region based on hydrogen-deuterium exchange in solution ^60^. Met98 connects with several residues from the 59–66 loop that in PTP1B includes a phosphorylation site (Tyr66) and was reported to be allosterically linked to active-site oxidation state ^61^, which is used for varying natural regulatory mechanisms in different PTPs ^62–65^. Leu260 is in the catalytic Q loop and connects with the P loop and α4 helix, which is allosterically linked to activity ^66,67^. Further down α4, Asp229 is at an allosteric activator site in STEP (PTPN5) ^34,68^, and connects with residues that exhibit conformational heterogeneity in high-resolution PTP1B structures ^67^ and enhance PTP1B activity when mutated ^66^. Finally, the 100% conserved Cys215 connects with several residues in the active-site P loop and E loop in our network, which is satisfying to observe given its catalytically essential nature. Overall, the residues with the most conserved dynamic interactions across PTPs are related to PTP catalysis and various modes of regulation.

To further dissect the sum network structure, we used the Girvan-Newman community detection algorithm ^69^ to partition the network, resulting in 7 communities or subnetworks (**Fig. 3c-d**). This suggests that the PTP fold is arranged in a hierarchical manner, with a small number of cohesive local communities or clusters that each experience collective dynamics internally. Some of these communities map to known functional regions, such as the catalytic Cys215 and nearby active-site residues (magenta) and the catalytic Q loop and substrate recognition loop (yellow).

### Network rewiring upon catalytic loop movement and ligand binding

We next sought to assess how the dynamic network common to all PTPs changes in concert with enzyme functional state. To do so, we compared subsets of structures with the catalytically critical, conformational bistable active-site WPD loop ^20,21,59^ in the closed state vs. open state (**Fig. 4a,d**), with an active-site ligand bound vs. the apo form (**Fig. 4b,d**), and with an allosteric ligand bound vs. the apo form (**Fig. 4c,d**). For each comparison, we ensured that resolution distributions were sufficiently similar for the subset networks (**Fig. S5**, **Fig. S6**, **Fig. S7**).

**Figure 4:**
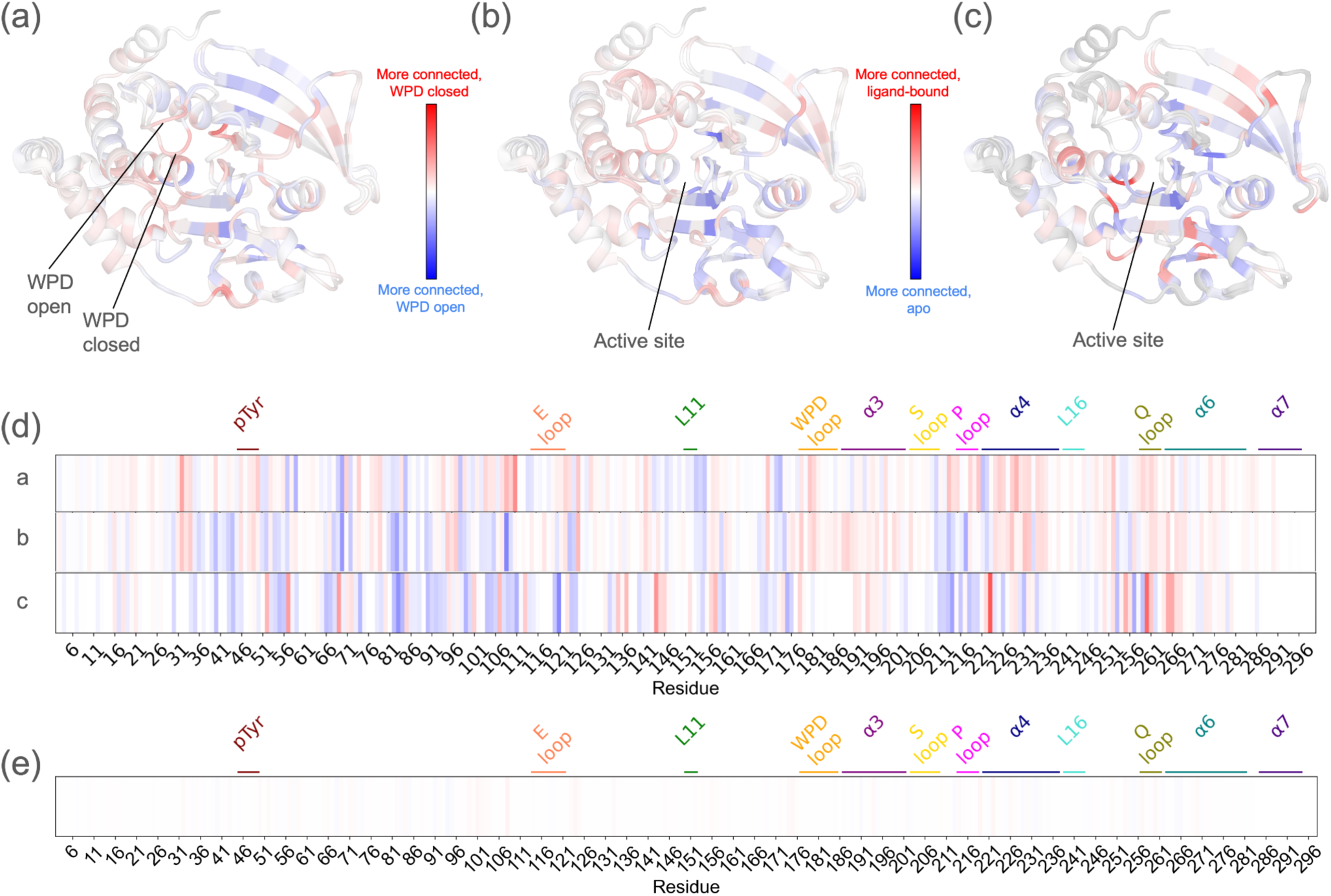
Rewiring of internal networks upon loop conformational change and ligand binding. The difference in weighted degree (Δdegree) for each residue in the all-PTPs sum network with all edges is mapped onto a cartoon visualization of structurally aligned, representative closed-state vs. open-state structures of the PTP catalytic domain (PDB ID: 1sug, 1t49) ^18,54^. See red/blue color bars. **(a)** WPD loop conformational changes. **(b)** Active-site ligand binding. **(c)** Allosteric ligand binding. **(d)** Δdegree from (a-c) is mapped onto a 1-dimensional representation of the protein sequence (PTP1B numbering), with key regions labeled. a/b/c labels on the left correspond to panels in the top row. **(e)** Δdegree is computed for 70 randomly sampled halves of our full dataset, averaged, and mapped onto a 1-dimensional representation. See also **Fig. S5**.

To assess changes in network connectivity, we mapped the difference in degree value (Δdegree) to the tertiary structure (**Fig. 4a-c**) and primary structure (**Fig. 4d**). For each comparison, degree changed substantially across the PTP catalytic domain, indicating dynamic rewiring of the structurally distributed internal network related to catalytic motions or ligand binding. Random sampling of different subsets of e.g. WPD closed vs. open structures leads to some variability but qualitatively similar Δdegree patterns (**Fig. S8**). By contrast, negative control calculations with randomly selected halves of all the structures in our dataset regardless of category yield an averaged Δdegree plot that is featureless (**Fig. 4e**).

When the WPD loop closes, degree increases moderately for several areas of the active site (red in **Fig. 4a,d**) including the WPD loop itself, P loop, Q loop, and pTyr binding loop. Degree also increases for other regions, including the Met109 region (see previous section) and allosteric α4 helix ^66,67^. This suggests that when they enter the closed “active” state, PTPs experience enhanced coupled conformational heterogeneity in the active site and related regions throughout the catalytic domain. At the same time, some other regions compensate with decreased coupled conformational heterogeneity (blue in **Fig. 4a,d**) including the allosteric Loop 11 (i.e. L11) ^21^.

Active-site (orthosteric) and non-active-site (allosteric) small-molecule ligands both induce significant Δdegree throughout the fold, but in different ways. The Δdegree pattern for active-site ligands is reminiscent of that for WPD loop closing (**Fig. 4a,d** vs. **Fig. 4b,d**) in that degree increases for the WPD loop, Q loop, and pTyr loop, yet degree decreases for the P loop, perhaps due to rigidification from the bound ligands. By contrast, the Δdegree pattern for allosteric ligands is distinct from that for WPD loop closing. This is likely because allosteric ligands bind at many locations (**Fig. S9**) that may have distinct effects on the network shared by all PTPs and/or on different tendrils of the network in different PTPs. Although there is a bias toward the WPD loop closed state for active-site ligands (62/80, 78%) and the open state for allosteric ligands (28/35, 80%), our control comparisons in the same WPD loop state also show different Δdegree patterns for active-site and allosteric ligands (**Fig. S7**), indicating these two ligand types impart fundamentally different dynamical effects on the PTP catalytic domain.

### Network rewiring between evolutionarily related PTPs

While the PTP family may share aspects of a consensus allosteric network ^35^, we hypothesized that this network has also been rewired in various ways for many PTPs over the course of evolution to diversify their regulation and function. To explore this hypothesis using RINFAIRE, we compared the sum network for each of several PTPs to the sum network for all other PTPs in our dataset. We selected PTP1B (**Fig. 5a,d**), SHP2 (**Fig. 5b,d**), and YopH (**Fig. 5c,d**) because they are the most abundant in our dataset (see **Fig. 1d** and Data availability - metadata), or in the case of YopH have been compared to PTP1B in previous studies ^59,70,71^, and have contributions from structures across a wide resolution range (**Fig. S10c**).

**Figure 5:**
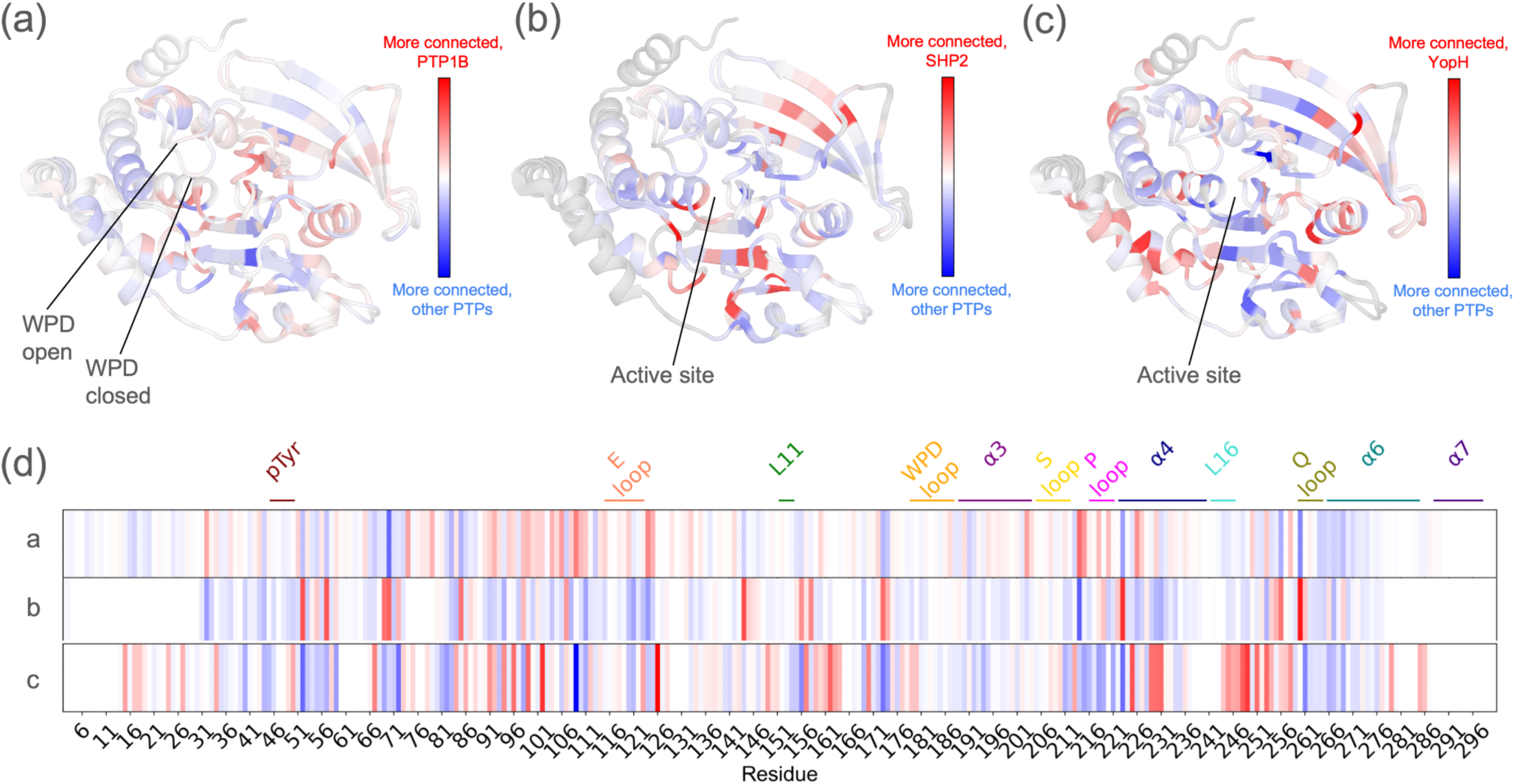
Rewiring of internal networks in specific PTPs within the PTP family. The difference in weighted degree (Δdegree) for each residue in the sum networks with all edges is mapped onto a cartoon visualization of structurally aligned, representative closed vs. open-state structures of the PTP catalytic domain (PDB ID: 1sug, 1t49) ^18,54^. See red/blue color bars. **(a)** PTP1B vs. other PTPs. **(b)** SHP2 vs. other PTPs. **(c)** YopH vs. other PTPs. **(d)** Δdegree is mapped onto a 1-dimensional representation of the protein sequence (PTP1B numbering), with key regions labeled. a/b/c labels on the left correspond to the panels in the top row. See also **Fig. S10** and **Fig. S11**.

The results reveal a distinct pattern of dynamic connectivity in each PTP (rows in **Fig. 5d**). PTP1B has the highest average Δdegree (+0.091), consistent with its well-known allosterism. SHP2 has the lowest average Δdegree (-1.067), likely because it is locked into the rigid autoinhibited open state in all known structures. YopH has an intermediate average Δdegree (+0.049). Its highest Δdegree regions correspond to the α4 helix, where mutations increase PTP1B activity ^66^, and the region surrounding D245, where a mutation decreases PTP1B activity ^72^. Because YopH is a highly active PTP, these observations suggest that changes in dynamics driven by sequence change in these regions of the PTP fold may play key roles in modulating catalytic activity.

The distribution of open vs. closed WPD states differs across PTPs, including PTP1B (37 vs. 41), SHP2 (35 vs. 0), and YopH (3 vs. 8). We therefore analyzed subsets of structures with the same WPD loop state, which resulted in similar Δdegree patterns (**Fig. S11**) as obtained from using all available structures (**Fig. 5d**). Together, these findings support our hypothesis that different PTPs exhibit distinct inherent allosteric wiring.

### Network overlap with residues involved in allostery/regulation/function

We next explored how the all-PTPs sum network from RINFAIRE overlapped with residues that were previously reported to be involved in allostery, dynamics, and/or other aspects of PTP function. We began by comparing our network to two so-called sectors of coevolving amino acid positions identified previously by statistical coupling analysis (SCA) for many PTP catalytic domain sequences ^35^. Sector A was associated with known allosteric regions, whereas the role of sector B was less well understood. In that work, residue positions with more nearby sector residues were associated with a higher fraction of experimentally characterized mutations that were functionally influential, based on a dataset of 67 experimentally characterized mutations spanning 13 PTPs. We performed the same analysis with our all-PTPs sum network, choosing an edge weight cutoff (top 3%) to closely match the combined size of both SCA sectors and thus maximize comparability. We observe a similar pattern, with mutations at sites near our network being more prone to influencing enzyme function (**Fig. 6a**). Specifically, only 43–52% of mutations at sites near 0–6 network residues influence function, yet 88–100% of mutations at sites near 6–12 network residues influence function (**Fig. 6a**).

**Figure 6:**
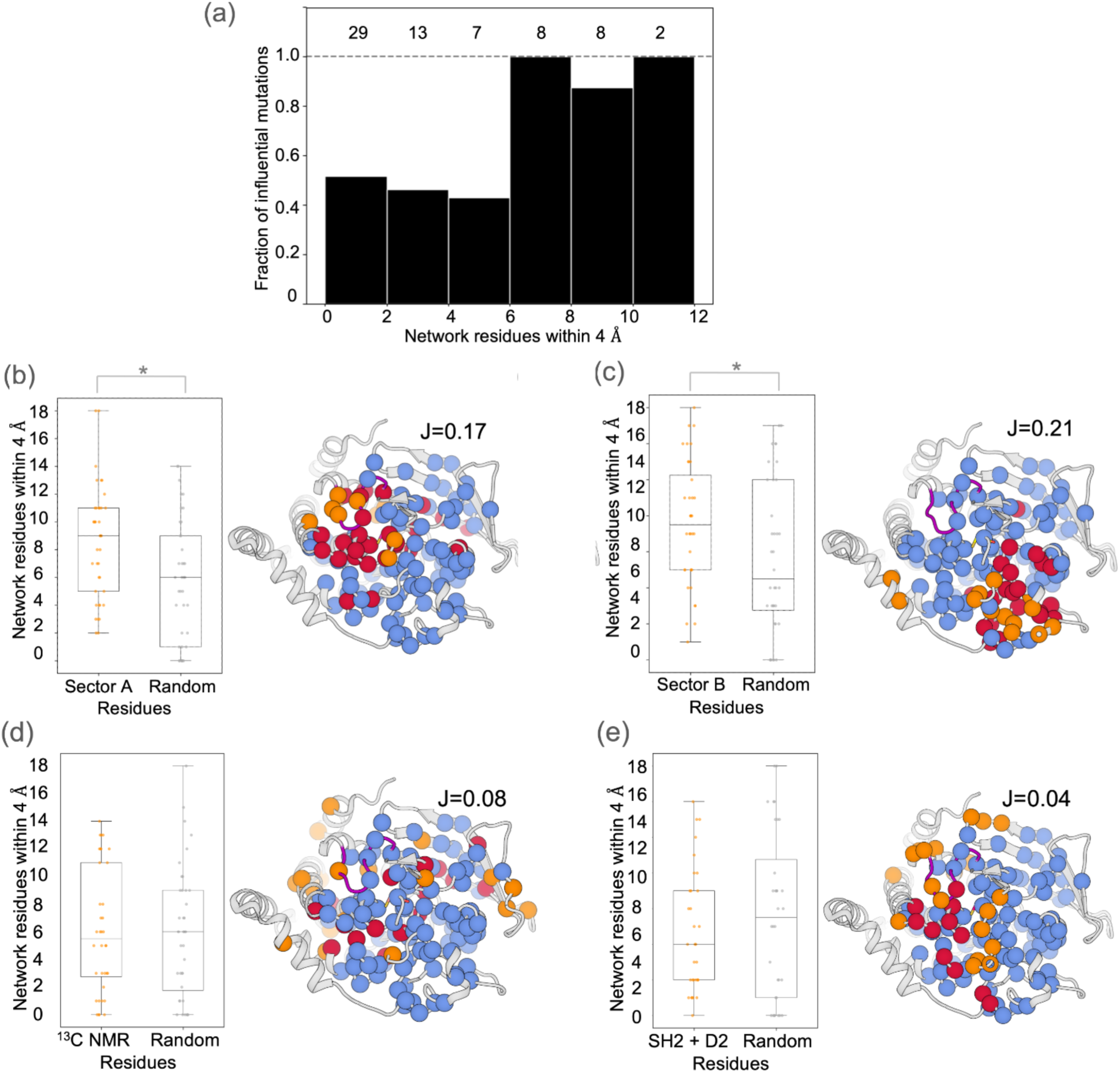
Colocalization of RINFAIRE network with regions of interest from previous studies. **(a)** Colocalization of our all-PTPs sum network (top 3% of edges) with previously experimentally characterized mutations ^35^. All residues in a representative structure of PTP1B (PDB ID: 3a5j) were binned based on the number of residues from our network nearby (x-axis). For each bin, all available curated experimentally characterized mutations (totals at top) were assessed, and the fraction that were functionally influential is indicated (y-axis). **(b-e)** Colocalization of our all-PTPs sum network (top 5% of edges) with different residues of interest from previous studies: **(b)** SCA sector A ^35^, **(c)** SCA sector B ^35^, **(d)** dynamic residues from ^13^C NMR for PTP1B ^73^, and **(e)** residues in regulatory domain interfaces with SH2 domains (SHP2) and D2 domains (receptor-type PTPs). *Left sub-panels:* Distribution of number of network residues within 4 Å for all residues in the set of interest, vs. similar analysis for random set of residues of the same size. * p < 0.05 indicates distributions are statistically significantly different from a Kolmogorov-Smirnov test. Jaccard ratio (J = intersection / union) is shown for each comparison between our network and residues of interest. *Right sub-panels:* Residues of interest (orange), our network residues (blue), and residues common to both (maroon) mapped to a representative structure of PTP1B (PDB ID: 1sug).

To examine the overlap of our network with the SCA sectors more directly, we used a statistical test that compared the number of nearby residues from our network for (i) a set of residues of interest relative to (ii) a random set of residues of the same size ^35^. The overlap was statistically significant both for our network with sector A and with sector B (**Fig. 6b-c**). Taken together, these results suggest that our dynamic structure-based network and the purely sequence-based sectors offer similar yet complementary insights into conserved allosteric wiring in the PTP catalytic domain.

We also explored how our network relates to sets of residues in the PTP fold that pertain to collective dynamics or specific modes of interdomain allosteric regulation. These include residues that exhibit intermediate-timescale dynamics from ^13^C NMR relaxation dispersion experiments for PTP1B ^73^ (**Fig. 6d**), or are located at regulatory domain interfaces with autoinhibitory SH2 domains in SHP2 or non-catalytic PTP-like D2 domains in receptor-type PTPs (**Fig. 1e**, **Fig. 6e**). In each case, the overlap with our network is not significant. However, there are caveats to these comparisons. First, ^13^C NMR experiments are limited to methyl-containing side chains and specific timescales, in contrast to our network which includes all atoms and is agnostic to timescales, and it is unknown to what extent similar dynamics exist in other PTPs beyond PTP1B. Second, the structural influences of regulatory domains may be felt beyond the direct interface residues that we chose to examine here; moreover, SHP2 operates by an autoinhibitory mechanism that is not present in other PTPs and may not necessitate allosteric signal propagation within the catalytic domain itself.

### Highly networked residues impact function regardless of sequence conservation

The preceding results suggest that the PTP network identified by RINFAIRE is relevant to allosteric modulation of enzyme activity (**Fig. 6a-c**). We experimentally tested this hypothesis in a forward manner by mutating residues implicated as being important in our network and characterizing their effects on enzyme activity. To identify suitable residues for these experiments, we examined the correlation between network weighted degree and sequence conservation across the PTP family. The correlation was moderate-to-weak (**Fig. 7a**), indicating that more conserved residues generally tend to be more dynamically interconnected, yet there is a range of connectivity for different residues within each bin of sequence conservation.

**Figure 7:**
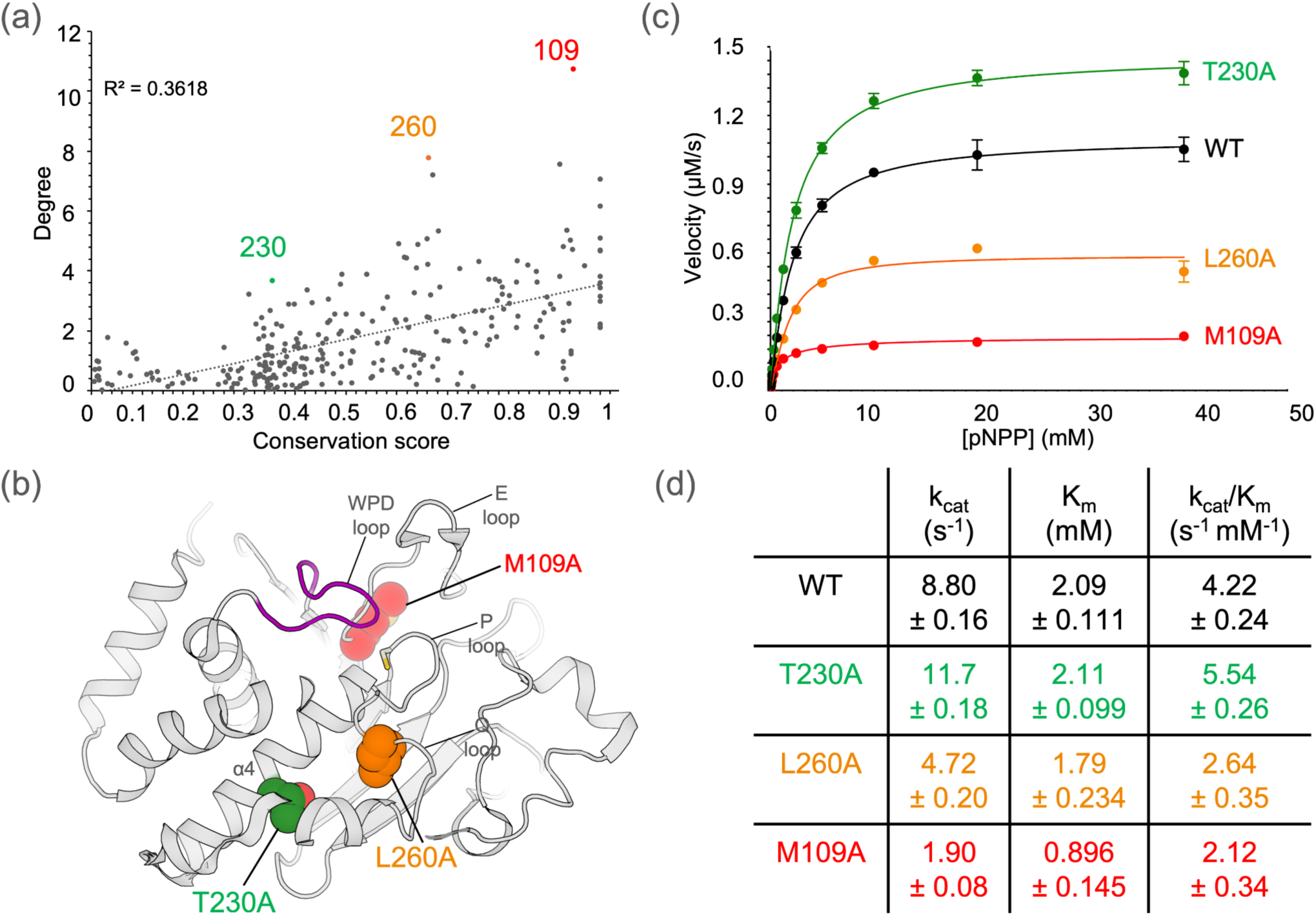
High-degree network residues control catalytic activity regardless of sequence conservation. **(a)** Weighted degree from our all-PTPs sum network plotted against sequence conservation, for all comparable residues in the PTP fold. Labeled, colored residues have high degree relative to their sequence conservation. R^2^ represents correlation for linear fit between degree values and conservation scores for all residues (dotted line). **(b)** Labeled residues from (a) are shown as spheres and mapped to a representative structure of PTP1B (PDB ID: 1sug), with colors corresponding to (a). Key regions including the WPD loop, E loop, P loop, Q loop, and α4 helix are labeled. **(c)** Experimental Michaelis-Menten kinetics plot using pNPP substrate for WT PTP1B vs. M109A, T230A, and L260A mutations. Data points represent average values from n=4 replicates; error bars represent 95% confidence intervals. **(d)** Michaelis-Menten kinetics parameters were derived from the average data in (c), with 95% confidence intervals indicating variability across replicates.

We therefore chose to mutate residues with high network connectivity given their sequence conservation, in three different conservation regimes: low (<40%), intermediate (40–80%), and high (>80%). These criteria led us to three promising, complementary residues: 230 (35.6% conserved), 260 (66.3%), and 109 (94.8%) (colored points in **Fig. 7a**). These residues are widely distributed in the 3D structure of PTP1B (**Fig. 7b**), but are all near functionally relevant sites, including the catalytic Cys215, catalytic Q loop, and allosteric α4 helix (**Fig. 7b**).

We subsequently created T230A, L260A, and M109A mutant proteins and performed enzyme activity assays (see Methods). Consistent with our hypothesis that these residues are integrally placed in the allosteric wiring of the PTP fold, all of these mutations significantly affect the catalytic activity of PTP1B significantly (**Fig. 7c-d**).

M109A reduces activity most dramatically, with a significant decrease in k_cat_ (∼4.6x). M109A also decreases K_m_ (∼2.3x), perhaps due to its proximity to the substrate-binding P loop. However, overall M109A significantly decreases k_cat_/K_m_ (∼2.0x). Our results for M109A are in line with prior reports that M109 mutations reduced activity by ∼8–10x ^35,74^. L260A reduces activity to an intermediate degree, with a decrease in k_cat_/K_m_ (∼1.6x) driven by a decrease in k_cat_ (∼1.9x).

Surprisingly, T230A, which is the most distal of the three mutations from the active site (**Fig. 7b**), enhances PTP1B activity, with an increase in k_cat_/K_m_ (∼1.3x, 30%) driven by an increase in k_cat_ (∼1.3x). Notably, several mutants of F225, which is roughly one turn away from T230 in the α4 helix, were also found to enhance activity in PTP1B, including F225Y (∼1.8x), F225Y/R199N (∼2.2x), and F225Y/R199N/L195R (∼4x) ^66^. These observations suggest that the broader α4 helix region in PTP1B, and potentially also in other PTPs ^66^, may play a central role in dictating the catalytic rate.

## Discussion

Despite a structurally conserved catalytic domain (**Fig. 1**), PTPs have divergent biological roles ^75,76^ that may be enabled by differences in allosteric wiring. Crystallographic multiconformer modeling with qFit ^45–49^ affords a unique opportunity to analyze coupling between alternate conformations that may underlie allostery, but methods to analyze these complex models have been limited. Here we introduce RINFAIRE, a new algorithm for analyzing networks of coupled conformational heterogeneity across related protein structures (**Fig. 2**). Coupling the latest improved version of qFit ^49^ to RINFAIRE, we have mapped a consensus PTP dynamic interaction network that encompasses many key catalytic and allosteric motifs (**Fig. 3**), analyzed how this network changes in response to catalytic motions and ligand binding (**Fig. 4**), assessed how it differs between functionally divergent PTPs (**Fig. 5**), compared it with various sets of dynamic/allosteric residues (**Fig. 6**), and validated it prospectively with *in vitro* biochemical experiments (**Fig. 7**). Together, our results suggest that the networks identified by RINFAIRE are indeed relevant to allostery in the PTP fold.

Future upstream developments of qFit could benefit downstream RINFAIRE analyses. First, qFit only models relatively small-scale alternate conformations (∼1 Å), so does not capture e.g. movements of the WPD loop, loop 16, and α7 helix ^21,67,77^ in PTPs. Future work can improve modeling of larger-scale backbone flexibility in qFit, e.g. using automated loop sampling driven by density maps ^78^ and/or cross-pollination of conformations from independent structures ^79^. Such modeling would be aided by new macromolecular model formats to encode hierarchical conformational heterogeneity ^80^. Second, small-molecule ligands bound to proteins can adopt alternate conformations in crystal structures ^47,48^, but qFit does not yet simultaneously model flexibility for both proteins and bound ligands. Future development can address this limitation, thus providing new opportunities to explore the interplay between protein and ligand conformational heterogeneity in e.g. active-site vs. allosteric-site binding pockets (**Fig. 4b,c**).

There is also room for future RINFAIRE developments that could yield new insights into mechanisms of allosteric wiring. First, RINFAIRE uses a distance-based approach (default: 4 Å) for identifying through-space residue-residue interactions and quantifying the associated edge weights (**Fig. 2b**). This approach has several advantages: simplicity, consistency with past precedent in the literature for protein structure RINs (albeit for static structures instead of alternate conformations) ^81–83^, and implicitly accounting for a variety of physicochemical interaction types including not only unfavorable steric clashes but also favorable H-bonds, van der Waals interactions, salt bridges, etc. These through-space interactions are complemented with through-backbone interactions (**Fig. 2b**), which also play important roles in correlated motions in proteins ^46,84,85^. Nevertheless, RINFAIRE is readily extensible to more complex/physics-based scoring functions for interactions between residues. Second, there is a growing algorithmic toolkit for protein structure contact network analysis that could prove useful for RINFAIRE, including modeling contact rearrangements as edges ^86^, eigenvector centrality for pinpointing allosteric residues ^87^, and many other ideas ^5,88,89^.

The analysis reported here benefits from the availability of many high-resolution crystal structures that sample distinct conformational states, crystal lattices, crystallization conditions, etc. and thus provide a useful “pseudo-ensemble” ^90–92^. For the PTP family, some PTPs are more well-represented in the PDB (**Fig. 1d**), which led us to focus our inter-PTP analyses on these PTPs (**Fig. 5**, **Fig. S11**). Careful matching of relevant experimental factors using the RINFAIRE metadata functionality may enable further inter-PTP comparisons which were beyond the scope of the current report. For example, specific crystal contacts may facilitate distinct patterns of local conformational states and/or disorder ^67,93^. Such comparisons will gain statistical power over time as more crystal structures are deposited to the PDB. Indeed, it is noteworthy that 153 of the 170 crystal structures used in this study were from the last 10 years. It is also possible that cryo-electron microscopy (cryo-EM) will reach the stage of yielding high-resolution structures for enzymes such as PTPs; notably, qFit also works with cryo-EM density maps ^49^. Additional alternative structures could be generated by computational means such as AlphaFold ^94,95^ with multiple sequence alignment subsampling ^96–98^, flow matching ^99^, or predicted side-chain χ angle distributions ^100^, and then used as inputs to RINFAIRE to predict allosteric networks at a larger scale, much as AlphaFold has been used at a proteome-wide scale ^101^.

Although we focused on the PTP enzyme family in this study, our new computational pipeline can be easily applied to any other sets of related protein structures with a sufficient number of suitable input structures. As such, it sets the stage for future studies of how conformational ensembles are reshaped by sequence changes to alter dynamic properties such as allosteric signaling in a variety of contexts, including other biomedically important protein families and trajectories of iteratively designed or ancestrally reconstructed proteins. Building on the ligand comparisons presented here (**Fig. 4b,c**), our pipeline could also be used to unveil allosteric effects of small-molecule fragment binding from high-throughput crystallographic screens ^21,77,102–104^, thus providing more confident footholds for rational allosteric drug design ^105^.

## Materials and Methods

The following is an abbreviated Materials and Methods section — for full details, see the Supplementary Information.

PTP catalytic domain structures were obtained using Pfam (PF00102) ^106^ and the PDB. Structures that were successfully automatically re-refined with PHENIX ^107–109^ were subjected to qFit multiconformer modeling ^49^, followed by removing non-catalytic domains and splitting individual catalytic domain instances in cases of non-crystallographic symmetry. Structure-based multiple sequence alignment was performed using PROMALS3D ^110^. Metadata including PTP name, crystallographic R-factors, ligand type and location, and WPD loop state were tabulated.

RINFAIRE generates a residue interaction network for each provided qFit multiconformer model based on spatial proximity of alternate conformations (within 4 Å); edges between residues are normalized based on residue size (number of atoms). Backbone alternate conformations of sequentially adjacent residues are treated differently with a recursive method. RINFAIRE then uses a multiple sequence alignment to construct a “multinetwork” with all residue numbers shifted to a common reference. In the multinetwork, all contributing networks (from individual structures) are log normalized based on their total edge weights, to discourage unbalanced contributions from networks with many connections (e.g. high-resolution structures). To generate a sum network, the total edge weight for each edge in the multinetwork is calculated. To facilitate most subsequent analyses, we trimmed the sum network to the top 5% of edges (95% of lowest edge weights removed).

To identify communities within the sum network, we used the Girvan-Newman method ^69^ implemented in NetworkX ^111^ and identified where modularity plateaus. For Δdegree plots, the degree values for all residues for two subset sum networks were subtracted, and the resulting differences visualized on the sequence and the structure with a common color scale. To ensure a comparable analysis across different datasets, one-tailed Mann-Whitney U tests were performed, and resolution ranges were adjusted as needed. Colocalization of the all-PTPs sum network with functionally influential experimentally characterized PTP mutations and statistical analysis of sum network overlap with other sets of residues of interest were performed as previously described ^35^. Residues in regulatory domain interfaces in SHP2 and D2-containing PTPs were identified using distance commands in PyMol.

PTP1B site-directed mutagenesis, expression, purification, and Michaelis-Menten enzyme activity assays were performed as previously described ^21,72^.

## Data availability

The following supplementary data files are available at this Zenodo repository:

https://doi.org/10.5281/zenodo.15420194.

- PTP structures metadata table.
- PTPs PROMALS3D multiple sequence alignment (MSA) file.
- Single-chain catalytic domain models from PTP qFit multiconformer structures (for use in all analyses).
- Full PTP qFit multiconformer structures (only for crystallographic refinement).
- Multinetwork Python pickle file for all-PTPs sum network with all edges.
- Residue weighted-degree values and residue-residue edge weights, for all-PTPs sum network with all edges (0% edges removed) and top 5% of edges (95% weakest edges removed).
- Lists of residues used for **Fig. 6**.

## Code availability

The open-source RINFAIRE software reported here is available at this GitHub repository: https://github.com/keedylab/rinfaire. The repository contains all Python code and scripts needed to run the software, a Pipfile to facilitate installation of dependencies, and a README file. The version used for the analyses in this study is v2025.1, the initial public release.

## Author contributions

**Akshay Raju:** conceptualization; methodology; investigation; software; formal analysis; visualization; writing – original draft; review and editing; data curation.

**Shivani Sharma:** conceptualization; methodology; investigation; formal analysis; visualization; writing – original draft; review and editing; data curation.

**Blake T. Riley:** conceptualization; data curation; investigation; methodology.

**Shakhriyor Djuraev:** investigation.

**Yingxian Tan:** data curation.

**Minyoung Kim:** data curation.

**Toufique Mahmud:** investigation.

**Daniel A. Keedy:** conceptualization; methodology; validation; supervision; writing – original draft; writing – review and editing; visualization; funding acquisition; resources; project administration.

## Acknowledgements

DAK is supported by NIH R35 GM133769. We thank Ali Ebrahim for help with structure visualization scripts, Virgil Woods for help with enzyme kinetics data analysis, and Stephanie Wankowicz and Henry van den Bedem for feedback on manuscript drafts.

## References

1. Monod, J. Chance and Necessity: An Essay on the Natural Philosophy of Modern Biology. Vintage Books (1971).

2. Fenton, A. W. Allostery: an illustrated definition for the ‘second secret of life’. Trends Biochem. Sci. 33, 420–425 (2008).

3. Gunasekaran, K., Ma, B. & Nussinov, R. Is allostery an intrinsic property of all dynamic proteins? Proteins 57, 433–443 (2004).

4. Swain, J. F. & Gierasch, L. M. The changing landscape of protein allostery. Curr. Opin. Struct. Biol. 16, 102–108 (2006).

5. Astore, M. A., Pradhan, A. S., Thiede, E. H. & Hanson, S. M. Protein dynamics underlying allosteric regulation. Curr. Opin. Struct. Biol. 84, 102768 (2024).

6. Papaleo, E. et al. The role of protein loops and linkers in conformational dynamics and allostery. Chem. Rev. 116, 6391–6423 (2016).

7. Daily, M. D. & Gray, J. J. Local motions in a benchmark of allosteric proteins. Proteins 67, 385–399 (2007).

8. van den Bedem, H., Bhabha, G., Yang, K., Wright, P. E. & Fraser, J. S. Automated identification of functional dynamic contact networks from X-ray crystallography. Nat. Methods 10, 896–902 (2013).

9. Popovych, N., Sun, S., Ebright, R. H. & Kalodimos, C. G. Dynamically driven protein allostery. Nat. Struct. Mol. Biol. 13, 831–838 (2006).

10. Lockless, S. W. & Ranganathan, R. Evolutionarily conserved pathways of energetic connectivity in protein families. Science 286, 295–299 (1999).

11. Lee, J. et al. Surface sites for engineering allosteric control in proteins. Science 322, 438–442 (2008).

12. Law, A. B., Fuentes, E. J. & Lee, A. L. Conservation of side-chain dynamics within a protein family. J. Am. Chem. Soc. 131, 6322–6323 (2009).

13. Campitelli, P., Modi, T., Kumar, S. & Ozkan, S. B. The role of conformational dynamics and allostery in modulating protein evolution. Annu. Rev. Biophys. 49, 267–288 (2020).

14. He, R.-J., Yu, Z.-H., Zhang, R.-Y. & Zhang, Z.-Y. Protein tyrosine phosphatases as potential therapeutic targets. Acta Pharmacol. Sin. 35, 1227–1246 (2014).

15. Tonks, N. K. Protein tyrosine phosphatases: from genes, to function, to disease. Nat. Rev. Mol. Cell Biol. 7, 833–846 (2006).

16. Alonso, A. et al. Protein tyrosine phosphatases in the human genome. Cell 117, 699–711 (2004).

17. Andersen, J. N. et al. Structural and evolutionary relationships among protein tyrosine phosphatase domains. Mol. Cell. Biol. 21, 7117–7136 (2001).

18. Wiesmann, C. et al. Allosteric inhibition of protein tyrosine phosphatase 1B. Nat. Struct. Mol. Biol. 11, 730–737 (2004).

19. Olmez, E. O. & Alakent, B. Alpha7 helix plays an important role in the conformational stability of PTP1B. J. Biomol. Struct. Dyn. 28, 675–693 (2011).

20. Choy, M. S. et al. Conformational Rigidity and Protein Dynamics at Distinct Timescales Regulate PTP1B Activity and Allostery. Mol. Cell 65, 644–658.e5 (2017).

21. Keedy, D. A., Hill, Z. B., Biel, J. T., Kang, E. & Rettenmaier, T. J. An expanded allosteric network in PTP1B by multitemperature crystallography, fragment screening, and covalent tethering. Elife (2018).

22. Singh, J. P. et al. Crystal Structure of TCPTP Unravels an Allosteric Regulatory Role of Helix α7 in Phosphatase Activity. Biochemistry 60, 3856–3867 (2021).

23. Xie, J. et al. Allosteric inhibitors of SHP2 with therapeutic potential for cancer treatment. J. Med. Chem. 60, 10205–10219 (2017).

24. Pádua, R. A. P. et al. Mechanism of activating mutations and allosteric drug inhibition of the phosphatase SHP2. Nat. Commun. 9, 4507 (2018).

25. van Vlimmeren, A. E. et al. The pathogenic T42A mutation in SHP2 rewires the interaction specificity of its N-terminal regulatory domain. Proc. Natl. Acad. Sci. U. S. A. 121, e2407159121 (2024).

26. Barr, A. J. et al. Large-scale structural analysis of the classical human protein tyrosine phosphatome. Cell 136, 352–363 (2009).

27. Wen, Y. et al. RPTPα phosphatase activity is allosterically regulated by the membrane-distal catalytic domain. J. Biol. Chem. 295, 4923–4936 (2020).

28. Elhassan, R. M., Hou, X. & Fang, H. Recent advances in the development of allosteric protein tyrosine phosphatase inhibitors for drug discovery. Med. Res. Rev. 42, 1064–1110 (2022).

29. Hansen, S. K. et al. Allosteric inhibition of PTP1B activity by selective modification of a non-active site cysteine residue. Biochemistry 44, 7704–7712 (2005).

30. Krishnan, N. et al. Targeting the disordered C terminus of PTP1B with an allosteric inhibitor. Nat. Chem. Biol. 10, 558–566 (2014).

31. Friedman, A. J. et al. Allosteric Inhibition of PTP1B by a Nonpolar Terpenoid. J. Phys. Chem. 126, 8427–8438 (2022).

32. Garcia Fortanet, J., et al. Allosteric inhibition of SHP2: Identification of a potent, selective, and orally efficacious phosphatase inhibitor. J. Med. Chem. 59, 7773–7782 (2016).

33. Chen, Y.-N. P. et al. Allosteric inhibition of SHP2 phosphatase inhibits cancers driven by receptor tyrosine kinases. Nature 535, 148–152 (2016).

34. Tautermann, C. S. et al. Allosteric activation of striatal-enriched protein tyrosine phosphatase (STEP, PTPN5) by a fragment-like molecule. J. Med. Chem. 62, 306–316 (2019).

35. Hjortness, M. K. et al. Evolutionarily Conserved Allosteric Communication in Protein Tyrosine Phosphatases. Biochemistry 57, 6443–6451 (2018).

36. Welsh, C. L. & Madan, L. K. Allostery in protein tyrosine phosphatases is enabled by divergent dynamics. J. Chem. Inf. Model. 64, 1331–1346 (2024).

37. Zhang, Z.-Y. Drugging the undruggable: Therapeutic potential of targeting protein tyrosine phosphatases. Acc. Chem. Res. 50, 122–129 (2017).

38. McClendon, C. L., Friedland, G., Mobley, D. L., Amirkhani, H. & Jacobson, M. P. Quantifying Correlations Between Allosteric Sites in Thermodynamic Ensembles. J. Chem. Theory Comput. 5, 2486–2502 (2009).

39. Supriyo, B. & Nagarajan, V. Differences in allosteric communication pipelines in the inactive and active states of a GPCR. Biophys. J. 107, 422–434 (2014).

40. LeVine, M. V. & Weinstein, H. NbIT--a new information theory-based analysis of allosteric mechanisms reveals residues that underlie function in the leucine transporter LeuT. PLoS Comput. Biol. 10, e1003603 (2014).

41. Singh, S. & Bowman, G. R. Quantifying allosteric communication via both concerted structural changes and conformational disorder with CARDS. J. Chem. Theory Comput. 13, 1509–1517 (2017).

42. Greener, J. G. & Sternberg, M. J. E. AlloPred: prediction of allosteric pockets on proteins using normal mode perturbation analysis. BMC Bioinformatics 16, 335 (2015).

43. Huang, W. et al. Allosite: a method for predicting allosteric sites. Bioinformatics 29, 2357–2359 (2013).

44. Cui, D. S., Beaumont, V., Ginther, P. S., Lipchock, J. M. & Loria, J. P. Leveraging Reciprocity to Identify and Characterize Unknown Allosteric Sites in Protein Tyrosine Phosphatases. J. Mol. Biol. 429, 2360–2372 (2017).

45. van den Bedem, H., Dhanik, A., Latombe, J. C. & Deacon, A. M. Modeling discrete heterogeneity in X-ray diffraction data by fitting multi-conformers. Acta Crystallogr. D Biol. Crystallogr. 65, 1107–1117 (2009).

46. Keedy, D. A. & Fraser, J. S. Exposing hidden alternative backbone conformations in X-ray crystallography using qFit. PLoS Comput. Biol. (2015).

47. van Zundert, G. C. P. et al. qFit-ligand Reveals Widespread Conformational Heterogeneity of Drug-Like Molecules in X-Ray Electron Density Maps. J. Med. Chem. 61, 11183–11198 (2018).

48. Riley, B. T. et al. qFit 3: Protein and ligand multiconformer modeling for X-ray crystallographic and single-particle cryo-EM density maps. Protein Sci. 30, 270–285 (2021).

49. Wankowicz, S. A. et al. Automated multiconformer model building for X-ray crystallography and cryo-EM. Elife 12 (2024).

50. Fenwick, R. B., van den Bedem, H., Fraser, J. S. & Wright, P. E. Integrated description of protein dynamics from room-temperature X-ray crystallography and NMR. Proc. Natl. Acad. Sci. U. S. A. 111, E445–54 (2014).

51. Wankowicz, S. A., de Oliveira, S. H., Hogan, D. W., van den Bedem, H. & Fraser, J. S. Ligand binding remodels protein side-chain conformational heterogeneity. Elife 11 (2022).

52. Brock, J. S. et al. A dynamic Asp-Arg interaction is essential for catalysis in microsomal prostaglandin E2 synthase. Proc. Natl. Acad. Sci. U. S. A. 113, 972–977 (2016).

53. wwPDB consortium. Protein Data Bank: the single global archive for 3D macromolecular structure data. Nucleic Acids Res. 47, D520–D528 (2019).

54. Iversen, L. F., Kastrup, J. S., Møller, K. B., Pedersen, A. K. & Peters, G. H. Water-molecule network and active-site flexibility of apo protein tyrosine phosphatase 1B. Acta Crystallogr. D Biol. Crystallogr. 60, 1527–1534 (2004).

55. Lisi, G. P. & Loria, J. P. Using NMR spectroscopy to elucidate the role of molecular motions in enzyme function. Prog. Nucl. Magn. Reson. Spectrosc. 92-93, 1–17 (2016).

56. Aricescu, A. R. et al. Structure of a tyrosine phosphatase adhesive interaction reveals a spacer-clamp mechanism. Science 317, 1217–1220 (2007).

57. Liu, S. et al. Targeting inactive enzyme conformation: aryl diketoacid derivatives as a new class of PTP1B inhibitors. J. Am. Chem. Soc. 130, 17075–17084 (2008).

58. Lang, P. T. et al. Automated electron-density sampling reveals widespread conformational polymorphism in proteins. Protein Sci. 19, 1420–1431 (2010).

59. Whittier, S. K., Hengge, A. C. & Loria, J. P. Conformational motions regulate phosphoryl transfer in related protein tyrosine phosphatases. Science 341, 899–903 (2013).

60. Woods, V. A., Abzalimov, R. R. & Keedy, D. A. Native dynamics and allosteric responses in PTP1B probed by high-resolution HDX-MS. Protein Sci. 33, e5024 (2024).

61. Saeed, A. A., Klureza, M. A. & Hekstra, D. R. Mapping protein conformational landscapes from crystallographic drug fragment screens. J. Chem. Inf. Model. 64, 8937–8951 (2024).

62. van Montfort, R. L. M., Congreve, M., Tisi, D., Carr, R. & Jhoti, H. Oxidation state of the active-site cysteine in protein tyrosine phosphatase 1B. Nature 423, 773–777 (2003).

63. Chiarugi, P. & Cirri, P. Redox regulation of protein tyrosine phosphatases during receptor tyrosine kinase signal transduction. Trends Biochem. Sci. 28, 509–514 (2003).

64. Tonks, N. K. Redox redux: revisiting PTPs and the control of cell signaling. Cell 121, 667–670 (2005).

65. Ostman, A., Frijhoff, J., Sandin, A. & Böhmer, F.-D. Regulation of protein tyrosine phosphatases by reversible oxidation. J. Biochem. 150, 345–356 (2011).

66. Torgeson, K. R. et al. Conserved conformational dynamics determine enzyme activity. Sci Adv 8, eabo5546 (2022).

67. Sharma, S., Skaist Mehlman, T., Sagabala, R. S., Boivin, B. & Keedy, D. A. High-resolution double vision of the allosteric phosphatase PTP1B. Acta Crystallogr. Sect. F Struct. Biol. Cryst. Commun. 80, 1–12 (2024).

68. Guerrero, L., et al. Three STEPs forward: A trio of unexpected structures of PTPN5. bioRxiv (2024) doi:10.1101/2024.11.20.624168.

69. Girvan, M. & Newman, M. E. J. Community structure in social and biological networks. Proc. Natl. Acad. Sci. U. S. A. 99, 7821–7826 (2002).

70. Brandão, T. A. S., Johnson, S. J. & Hengge, A. C. The molecular details of WPD-loop movement differ in the protein-tyrosine phosphatases YopH and PTP1B. Arch. Biochem. Biophys. 525, 53–59 (2012).

71. Moise, G. et al. A YopH PTP1B chimera shows the importance of the WPD-loop sequence to the activity, structure, and dynamics of protein tyrosine phosphatases. Biochemistry 57, 5315–5326 (2018).

72. Perdikari, A. et al. Structures of human PTP1B variants reveal allosteric sites to target for weight loss therapy. bioRxiv (2024) doi: 10.1101/2024.08.05.603709.

73. Torgeson, K. R., Clarkson, M. W., Kumar, G. S., Page, R. & Peti, W. Cooperative dynamics across distinct structural elements regulate PTP1B activity. J. Biol. Chem. 295, 13829–13837 (2020).

74. Hjortness, M. K. et al. Abietane-type diterpenoids inhibit protein tyrosine phosphatases by stabilizing an inactive enzyme conformation. Biochemistry 57, 5886–5896 (2018).

75. Tabernero, L., Aricescu, A. R., Jones, E. Y. & Szedlacsek, S. E. Protein tyrosine phosphatases: structure-function relationships. FEBS J. 275, 867–882 (2008).

76. Tonks, N. K. & Neel, B. G. From form to function: signaling by protein tyrosine phosphatases. Cell 87, 365–368 (1996).

77. Skaist Mehlman, T., et al. Room-temperature crystallography reveals altered binding of small-molecule fragments to PTP1B. Elife 12 (2023).

78. van den Bedem, H., Lotan, I., Latombe, J. C. & Deacon, A. M. Real-space protein-model completion: an inverse-kinematics approach. Acta Crystallogr. D Biol. Crystallogr. 61, 2–13 (2005).

79. Keedy, D. A. Journey to the center of the protein: allostery from multitemperature multiconformer X-ray crystallography. Acta Crystallogr D Struct Biol 75, 123–137 (2019).

80. Wankowicz, S. A. & Fraser, J. S. Comprehensive encoding of conformational and compositional protein structural ensembles through the mmCIF data structure. IUCrJ 11, 494–501 (2024).

81. Salamanca Viloria, J., Allega, M. F., Lambrughi, M. & Papaleo, E. An optimal distance cutoff for contact-based Protein Structure Networks using side-chain centers of mass. Sci. Rep. 7, 2838 (2017).

82. Yuan, C., Chen, H. & Kihara, D. Effective inter-residue contact definitions for accurate protein fold recognition. BMC Bioinformatics 13, 292 (2012).

83. Yao, X.-Q., Momin, M. & Hamelberg, D. Establishing a Framework of Using Residue-Residue Interactions in Protein Difference Network Analysis. J. Chem. Inf. Model. 59, 3222–3228 (2019).

84. Davis, I. W., Arendall, W. B., 3rd, Richardson, D. C. & Richardson, J. S. The backrub motion: how protein backbone shrugs when a sidechain dances. Structure 14, 265–274 (2006).

85. Fenwick, R. B., Orellana, L., Esteban-Martín, S., Orozco, M. & Salvatella, X. Correlated motions are a fundamental property of β-sheets. Nat. Commun. 5, 4070 (2014).

86. Daily, M. D., Upadhyaya, T. J. & Gray, J. J. Contact rearrangements form coupled networks from local motions in allosteric proteins. Proteins 71, 455–466 (2008).

87. Negre, C. F. A. et al. Eigenvector centrality for characterization of protein allosteric pathways. Proc. Natl. Acad. Sci. U. S. A. 115, E12201–E12208 (2018).

88. Feher, V. A., Durrant, J. D., Van Wart, A. T. & Amaro, R. E. Computational approaches to mapping allosteric pathways. Curr. Opin. Struct. Biol. 25, 98–103 (2014).

89. Di Paola, L. & Giuliani, A. Protein contact network topology: a natural language for allostery. Curr. Opin. Struct. Biol. 31, 43–48 (2015).

90. Best, R. B., Lindorff-Larsen, K., DePristo, M. A. & Vendruscolo, M. Relation between native ensembles and experimental structures of proteins. Proc. Natl. Acad. Sci. U. S. A. 103, 10901–10906 (2006).

91. Yabukarski, F. et al. Assessment of enzyme active site positioning and tests of catalytic mechanisms through X-ray-derived conformational ensembles. Proc. Natl. Acad. Sci. U. S. A. 117, 33204–33215 (2020).

92. Yabukarski, F. et al. Ensemble-function relationships to dissect mechanisms of enzyme catalysis. Sci. Adv. 8, eabn7738 (2022).

93. Tyka, M. D. et al. Alternate states of proteins revealed by detailed energy landscape mapping. J. Mol. Biol. 405, 607–618 (2011).

94. Jumper, J. et al. Highly accurate protein structure prediction with AlphaFold. Nature 596, 583–589 (2021).

95. Abramson, J. et al. Accurate structure prediction of biomolecular interactions with AlphaFold 3. Nature 630, 493–500 (2024).

96. Del Alamo, D., Sala, D., Mchaourab, H. S. & Meiler, J. Sampling alternative conformational states of transporters and receptors with AlphaFold2. Elife 11, e75751 (2022).

97. Wayment-Steele, H. K. et al. Predicting multiple conformations via sequence clustering and AlphaFold2. Nature 625, 832–839 (2024).

98. Monteiro da Silva, G., Cui, J. Y., Dalgarno, D. C., Lisi, G. P. & Rubenstein, B. M. High-throughput prediction of protein conformational distributions with subsampled AlphaFold2. Nat. Commun. 15, 2464 (2024).

99. Jing, B., Berger, B. & Jaakkola, T. AlphaFold meets flow matching for generating protein ensembles. arXiv (2024) doi: 10.48550/arXiv.2402.04845.

100. Cagiada, M., Thomasen, F. E., Ovchinnikov, S., Deane, C. M. & Lindorff-Larsen, K. AF2χ: Predicting protein side-chain rotamer distributions with AlphaFold2. bioRxiv (2025) doi: 10.1101/2025.04.16.649219.

101. Varadi, M. et al. AlphaFold Protein Structure Database: massively expanding the structural coverage of protein-sequence space with high-accuracy models. Nucleic Acids Res. 50, D439–D444 (2022).

102. Douangamath, A. et al. Crystallographic and electrophilic fragment screening of the SARS-CoV-2 main protease. Nat. Commun. 11, 5047 (2020).

103. Schuller, M. et al. Fragment binding to the Nsp3 macrodomain of SARS-CoV-2 identified through crystallographic screening and computational docking. Sci Adv 7, (2021).

104. Mehlman, T., Ginn, H. M. & Keedy, D. A. An expanded trove of fragment-bound structures for the allosteric enzyme PTP1B from computational reanalysis of large-scale crystallographic data. Structure (2024) doi:10.1016/j.str.2024.05.010.

105. Krojer, T., Fraser, J. S. & von Delft, F. Discovery of allosteric binding sites by crystallographic fragment screening. Curr. Opin. Struct. Biol. 65, 209–216 (2020).

