## Supplementary Methods and Figures for "Mapping allosteric rewiring in related protein structures from collections of crystallographic multiconformer models"

#### Supplementary Materials and Methods

##### Dataset curation and processing

To obtain a list of structures that mapped to the "classical" protein tyrosine phosphatase family found in humans and other homologs, we used the Pfam protein family PF00102 [106] as an initial grouping of sequences and structures. Only natural PTP enzymes were included; engineered chimeric PTPs [112,113] were excluded. Similarly, archaeal PTPs with a sequence similarity of <30% were excluded [114]. Structures with only a catalytically inactive D2 domain (no D1 domain) were excluded as well. These entries were then filtered to only include those with protein structures in the Protein Data Bank (PDB) (as of June 17th, 2022). Two additional structures were manually added into our analysis, for TCPTP (PTPN2) (PDB ID: 7f5n and 7f5o). The structures (.pdb format) and structure factor data (.mtz format) were then inputted into a Jupyter Notebook, where they were processed using the Python package GEMMI [115]. Structures were then filtered to only those resolved using X-ray crystallography and having a resolution equal to or better than 2.1 Å, leaving a total of 189 structures for further analysis.

Automated structure refinement was performed through the PHENIX software (version 1.19.2-4158) [107–109]. To prepare the model for refinement, phenix.ready_set was run to add hydrogens and create .cif restraints files for any ligands. For refinement, phenix.refine was then run using the following parameters:

- *.pdb file from phenix.ready_set*
- *.mtz file*
- *.cif restraints file(s) for ligand(s) generated by phenix.ready_set*
- *refinement.refine.strategy=individual_sites+individual_adp+occupancies*
- *refinement.main.nqh_flips=true*
- *optimize_xyz_weight=true*
- *optimize_adp_weight=true*
- *hydrogens.refine=riding*
- *refinement.main.number_of_macro_cycles=8*
- *refinement.output.write_def_file=false*
- *refinement.output.write_eff_file=false*
- *refinement.output.write_geo_file=false*
- *refinement.input.xray_data.labels={xraylabel}*
- *refinement.input.xray_data.r_free_flags.{rfreelabel}*

The X-ray label and $R_{free}$ label were determined from the output of the phenix.mtz.dump utility. To test whether the refinement had in fact improved the structure to better fit the data, the $R_{free}$, $R_{work}$, and R-gap ($R_{free}$ - $R_{work}$) values before and after the refinement were then aggregated and compared, with structures that had an increase of ≥2.5% in $R_{free}$ ($R_{free(start)}$ - $R_{free(final)}$) after refinement being removed (**Fig. S12**). In addition, 8 structures failed during automated re-refinement, and were excluded from further analysis. Composite omit maps for input to qFit were generated using phenix.composite_omit_map.

#### Multiconformer modeling with qFit

We used qFit to identify alternate conformations of proteins that are supported by the electron density but were not initially modeled. Briefly, qFit samples possible conformations of each residue's backbone and sidechain, and selects the set of discrete alternate conformations that best and most parsimoniously explain the local electron density. It then reassembles the protein, including flexible backbone segments, to generate a complete but unrefined multiconformer model of the protein. This model is then refined using PHENIX and low-occupancy conformations are iteratively culled, yielding the final qFit multiconformer model. Previous versions of qFit introduced the algorithm [45], added backbone flexibility [46], added support for small-molecule ligands [47], added support for cryo-EM as well as X-ray maps [48], and provided further algorithmic enhancements [49].

For this study, we used a version (untagged) slightly ahead of the most recently released qFit version 3.2.2, with additional development (until and including commit #372) [49]. We used the following command line parameters for qFit:

```
qfit_protein \
composite_omit_map.mtz
-l 2FOFCWT,PH2FOFCWT \
refined_structure.pdb \
-d output_directory \
-p 20
```

For the final iterative refinement stage, we used the built-in qFit refinement script with PHENIX (version 1.19.2-4158). Of the 189 input structures, 6 were excluded due to an increase in $R_{free}$ ≥ 2.5% during initial rounds of refinement. An additional 6 structures failed during qFit multiconformer modeling and/or the final iterative refinement stage. Furthermore, 7 structures exhibited an $R_{free}$ increase of ≥ 2.5% during the qFit final refinement. These 19 structures were excluded from further analysis, resulting in a final dataset of 170 structures. These intermediate, full-asymmetric-unit qFit models are available as supplementary information.

The PTP qFit structures were then processed to remove extra protein domains within the same polypeptide chain, such as regulatory SH2 domains and non-catalytic D2 domains, so that the analysis would be confined to just the PTP catalytic domain. Non-catalytic D2 domains, which are in the same polypeptide as the main catalytic D1 domain, were not considered to be additional catalytic domain structures for our study. Some structures contained multiple non-identical copies of the catalytic domain by non-crystallographic symmetry; these domains were split into separate model files for subsequent analysis.  These final catalytic domain qFit models are available as supplementary information.

#### Multiple sequence alignment and metadata

To prepare inputs for the RINFAIRE program (see below), PROMALS3D [110] was used to generate a structure-based multiple sequence alignment (MSA) for all the structures in the dataset .  The output MSA was used to calculate the conservation score per residue using the ScoreCons server [116].  The PTPs MSA file is available as supplementary information.

We curated a metadata table using information about PTP crystal structures deposited in the PDB. We collected data on source organism, protein name, gene name, resolution, R-factors, ligand status (bound vs. apo), nature of ligand (inhibitory vs. activating vs. no effect), ligand binding location (active vs. allosteric), mutations (if any), and domains modeled.  Visual inspection in PyMol [117] was used to also identify the state of the WPD loop (open, closed, or super-open).  The metadata table is available as supplementary information.

To calculate the average sequence identity in the catalytic domain across classical human PTPs (**Fig. 7a**), Clustal Omega [118] was used to perform a multiple sequence (MSA) alignment of the wild-type sequence for all 37 human PTPs, using only the catalytic domain.  The resulting sequenced-based MSA is distinct from the structure-based MSA used for aligning networks within RINFAIRE (see below).  Mean sequence identity values were calculated using the sequence-based MSA alignment matrix result file.

#### Overview of the core RINFAIRE program

##### Constructing individual networks

RINFAIRE takes in a set of multiconformer protein structures (.pdb files) and a sequence alignment of the protein sequences (in the case that there is more than one structure).  The program starts by generating individual networks of conformationally coupled residues in each input protein structure. These networks are undirected weighted graphs in which the nodes represent residues and the edges represent the conformational coupling between residues.  To find the degree to which two residues' sets of alternate conformations are conformationally linked, we employ a distance-based approach, while treating alternate conformations along the backbone of two consecutive residues differently.

For every pair of residues, RINFAIRE identifies the atoms that have an alternate conformation in the structure, including hydrogen atoms added to the model.  For residues that are not sequentially adjacent, it first finds all pairs of alternate conformations between the two residues.  Because the alternate-location (alt-loc) labels in both residues might not reflect how they are coupled, we search across all possible pairs including those with the same label.  For each pair of conformers, it calculates the distances for all atoms between them and sums the number of atoms that are within 4 Å of each other.  While this parameter is adjustable in our program, we chose a 4 Å cutoff distance because previous literature had suggested that distance thresholds around 4 Å are a reasonable cutoff point for residue-residue contact analysis [83].

We then take the sum of all atom counts for all pairs of conformations for a given residue pair and normalize this count by the total number of atoms across all conformers in both residues. This is to mitigate biases both from larger residues having more possible connections along with residues that have many alternate conformations that might also inflate the number of connections between residues. The normalized count represents the combined measure of connectivity between these two residues. If this value is not zero, an edge is drawn between the two residues in the network with a weight equal to the normalized count.

Pairs of residues that are sequentially adjacent are treated differently, as there could be interactions along the backbone as well as steric interactions between both the backbone and sidechain and between the two sidechains themselves. To model backbone-backbone interactions we use an algorithm that progressively searches for alternate conformations across the backbone of two residues and tallies the number of alternate conformation atoms along each path. In this method, we only iterate over the same alt-loc label across both residues (ie. alt A of residue x and alt A of residue x+1). For each pair, it starts by checking if the atoms across the amide bond between the two residues have alternate conformations, since any backbone movement between the two residues must pass through this. If so, then it recursively searches the next chemically bonded atom along the backbone and adds that to the count of atom connections if that is also an alternate conformation for that alt-loc label.

Sidechain-sidechain and sidechain-backbone connections are calculated using the distance based algorithm with the same 4 Å cutoff metric. Due to the proximity of the beta carbon on the sidechain to the rest of the backbone, we removed the beta carbon along with all of the hydrogens bonded to it when considering sidechain-backbone steric interactions. Once all of the backbone-backbone, sidechain-backbone, and sidechain-sidechain interactions are counted and normalized, this value is the total connectivity between the two residues and added as an edge in the network.

##### Constructing the multinetwork

Once these networks are created for each individual structure, we then align each structure's network using the user-provided sequence alignment. This is done by shifting the residue number for each individual network residue to the corresponding position of that residue in the alignment. This allows for analogous residues across structures to have node labels that map onto the same alignment position even if the two structures are homologs, have unmodeled regions, or have different residue numbering schemes. This shifting can always be undone in later stages of the pipeline when we need to map alignment residue position back onto a reference structure's position by using the sequence alignment.

The network corresponding to each structure is log normalized based on the log of the total edge weight across the individual network relative to other individual networks. This normalization is intended to put networks with different numbers of edges, which may stem from structures with different numbers of alternate conformations (due to factors such as differing crystallographic resolution), on comparable footing. After this transformation, the total edge weights are also clipped at the 99th percentile of the distribution of total edge weights across all structures so that any outlier structures with a much larger number of edges do not overly skew the overall network.

Aligning the individual networks allows us to easily compare them at analogous residue positions. Internally, since we can represent each shifted network as an adjacency matrix, we can simply stack each n x n adjacency matrix (where n is the length of the sequence alignment) on top of each other. This creates an n x n x m dimensional array (where m is the number of structures) that we call the multinetwork. This object is what then gets passed to downstream analyses that will take the sum, subset, and perform other operations on this data.

#### Sum network analysis

The sum network was generated by using the aligned multinetwork object and taking the sum of the edge weights across all structures in the dataset. This was achieved by taking the sum across the structure dimension of the multinetwork array such that we get an n x n matrix that is also an adjacency matrix of the summed network for the entire dataset. Unless otherwise noted for some analyses, we then removed 95% of the weakest edges by edge weight, and removed any component network with less than five residues. While these parameters resulted in easily interpretable networks for our system, we allow these values to be altered by the user. Finally, the network was also shifted back to the reference sequence of PTP1B (PDB ID: 1sug) at analogous positions on the sequence alignment.

To identify communities within the sum network, we used the Girvan-Newman method for community detection [69] implemented in the Python library NetworkX [111]. The modularity score for each number of partitions was calculated, with the best partition being picked when the increase in modularity score had plateaued (increase from k partitions to k+1 partitions was < 0.01) (**Fig. S4**).

An additional consideration when analyzing sum networks concerns the WPD loop, loop 16 (L16), and α7 helix (in PTP1B and TCPTP). Although these regions are highly dynamic and critical to PTP function, the list of most connected (highest-degree) residues excludes them. This is likely because the WPD loop and loop 16 open/closed movements are large (each ~6 Å) and α7 undergoes an order-disorder transition, neither of which can be automatically modeled by qFit currently. As a result, these regions are not modeled with crystallographic alternate conformations (with relatively rare exceptions in the PDB [21,77,119,120]), so their importance is not captured by RINFAIRE.

#### Degree difference plots

A pair of sum networks were used to calculate the difference in degree per residue. At the time of running analysis_sum.py, -–seq_to_ref flag was used with a single reference structure (PDB: 1sug, chain A) to keep the residue numbering consistent for downstream difference calculations. For visualization, an RGB spectrum was used with an absolute color scale for consistency across comparisons. The absolute maximum Δdegree for each plot was set at a value of 10 and used for all the analysis. The same scale is used for visualizing as 1-dimensional strip plots and as 3-dimensional structure cartoons with PyMol. The same steps were used for comparing random subsets of structures (**Fig. S8**).

We carried out multiple comparative analyses using the sum networks from different PTPs as well as the different states in PTPs. The output sum network for each condition was generated using RINFAIRE. The degree value for every residue in the network was then used to calculate the difference, comparing the two datasets. This includes subsets of PTP sum networks, based on the state of their WPD loop (open vs. closed), ligand state (bound vs. apo), and individual PTPs such as PTP1B, SHP2, and YopH, each compared to all other PTPs. To ensure that comparable sets of structures were being used for each comparison, a one-tailed Mann-Whitney U test was performed to compare the resolution distribution for the structures. This is a suitable test for our data because it is non-parametric and does not assume a normal distribution (**Fig. S5**-**7, Fig. S10**).

#### Defining regulatory interface in SHP2 and D1/D2 structures

All structures in our analysis with an SH2 domain or D2 domain were used to calculate interface residues for SHP2 structure and D2-domain-containing structures respectively. The distance cutoff for the interface was set at 4 Å and each domain was defined for calculation. For SH2 domains, the PyMol command used to obtain interface residues was: '*select near_SH2, (byres *_* and i. 225-517) within 4 of (*_* and i. 1-215)*'. This resulted in a list of residues including (SHP2 numbering) 229, 244, 248, 249, 252, 253, 255, 256, 257, 258, 259, 260, 262, 265, 279, 280, 281, 282, 285, 364, 366, 425, 426, 427, 460, 461, 463, 464, 465, 502, 503, 506, 507, 508, and 510. For D2 domains, the PyMol commands to select interface residues were: '*select D1, (2FH7 or 4BPC) and i. 1368-1650*'; '*sele D2, (2FH7 or 4BPC) and i. 1659-1942*'; '*sele D1_near_D2, byres (D1 within 4 of D2)*'. This resulted in a list of residues including (D1/D2 numbering) 1526, 1527, 1562, 1563, 1565, 1566, 1572, 1573, 1647, 1650, and 1525.

#### Network overlap analysis

The sum network for colocalization analysis (**Fig. 6a**) was constructed using a slightly different edge weight cutoff (removing 97% of the weakest edges) from most other analyses, which resulted in a total of 82 residues. This was to approximately match the combined size of both SCA sectors of 75 residues [35]. By contrast, our default edge weight cutoff (removing 95% of edges) has a total of 88 residues. The lists of "influential" and "experimentally characterized" mutations were compiled from previous literature [35]. Each bin is inclusive of the lower bound but excludes the upper bound; thus the first bin (0–2) includes residues that are not within ≤ 4 Å of any residues from our network. The fraction was calculated using the number of residues in the influential mutation category in each bin divided by the number of residues in that bin that have been experimentally characterized.

For the overlap analyses with different sets of residues of interest (**Fig. 6b-e**), a Kolmogorov-Smirnov non-parametric test was used to measure statistical significance. This overlap is assessed between the region of interest and either the set of highly connected residues in our network (top 5% edges) or a set of randomly selected network residues (no edges removed). The latter analysis was repeated 100 times, each using a different randomly selected set of residues. The sampling shown in the main figure (**Fig. 6b–e**) corresponds to the final random sample, which we confirmed yields a p-value that is consistent with the majority of the samples ($p < 0.05$ for 89/100 in **Fig. 6b**, 76/100 in **Fig. 6c**, 0/100 in

**Fig. 6d**, and 2/100 in **Fig. 6e**) and thus is representative of the overall distribution of random samples. The use of the KS test for such analyses has precedent in prior literature [35].

#### Enzyme expression and purification

All biophysical experiments were performed using the wild-type PTP1B sequence comprising residues 1–321. The construct was cloned into a pET24b vector, which includes a kanamycin resistance gene. Unlike some previous crystallographic studies involving PTP1B, this work utilized the true wild-type sequence, without the commonly used WT* mutations (C32S/C92V). The initial wild-type construct contained residues 1–435 of PTP1B, but site-directed mutagenesis was previously employed to truncate it to residues 1–321. Using this shortened construct as a template, site-directed mutagenesis was also applied to generate the M109A, T230A, and L260A variants.

Protein expression and purification followed a previously established protocol with minor modifications. Plasmids carrying the intended mutations were introduced into competent *E. coli* BL21 (DE3) cells via transformation. After overnight incubation on LB agar plates supplemented with kanamycin at 37°C, individual colonies were used to inoculate 5 mL LB cultures containing kanamycin (1 mM final concentration), which were grown overnight at 37°C with shaking. The overnight cultures were then used to inoculate larger 1 L LB cultures with the same antibiotic concentration. These were grown at 37°C with shaking until the optical density at 600 nm ($OD_{600}$) reached approximately 0.6–0.8. Protein expression was induced with IPTG at a final concentration of 500 µM, and cultures were incubated overnight at 18°C with shaking. The cells were collected via centrifugation, flash-frozen, and stored at -80°C in 50 mL conical tubes until further purification.

For purification, cell pellets ("cellets") were resuspended in a lysis buffer containing Pierce protease inhibitor tablets and vortexed thoroughly. The suspension was sonicated on ice for 10 minutes at 50% amplitude, using 10-second on/off pulses. Following sonication, the lysate was centrifuged, and the supernatant was filtered through a 0.22 µm syringe filter before proceeding with purification. The first purification step involved cation exchange chromatography using a HiPrep SP FF 16/10 column (GE Healthcare Life Sciences), with a lysis buffer containing 100 mM MES (pH 6.5), 1 mM EDTA, and 1 mM DTT, alongside a NaCl gradient ranging from 0 to 1 M. The target protein eluted at approximately 200 mM NaCl. This was followed by size exclusion chromatography on an S75 column (GE Healthcare Life Sciences) using a buffer composed of 10 mM Tris (pH 7.5), 0.2 mM EDTA, 25 mM NaCl, and 3 mM DTT. The purity of the final protein sample was confirmed through SDS-PAGE analysis, which indicated a high level of purity with no detectable contaminants.

#### Enzyme activity assays

To assess the kinetic parameters of the mutant proteins, a colorimetric assay was performed using *para*-nitrophenyl phosphate (pNPP) as the substrate. The assay buffer was prepared with a final composition of 50 mM HEPES (pH 7.0), 1 mM EDTA, 100 mM NaCl, 0.05% Tween-20, and 1 mM β-mercaptoethanol (BME). After being filtered through a 0.22 µm membrane, the buffer was stored at room temperature. A series of 12 pNPP concentrations, ranging from 40 mM to 0.039mM, was generated via serial dilution in the assay buffer to ensure a wide range of substrate concentrations for kinetic analysis.

Before initiating the assay, the concentration of each mutant protein was measured twice in three independent replicates using a NanoDrop One. The protein samples were then diluted to a uniform concentration of 125 nM in the assay buffer, and the final concentration of each mutant protein was re-evaluated to confirm consistency. For the assay, 50 µL of the diluted protein solution was dispensed into wells of a Corning 96-well flat-bottom, non-binding polystyrene plate. The reaction was initiated by adding 50 µL of the pNPP + assay buffer solution to each well, followed by gentle pipetting to ensure thorough mixing. Absorbance at 405 nm was recorded every 17 seconds over a 6-minute period using a SpectraMax i3 plate reader. Each pNPP concentration was tested in quadruplicate for each mutant protein.

The rate of absorbance change (mAU per minute) over the 6-minute duration was determined and used to calculate the maximum reaction velocity ($V_{max}$). The catalytic constant ($k_{cat}$) was obtained by dividing $V_{max}$ by the average concentration of the corresponding mutant protein. Kinetic values from two independent experiments were pooled and analyzed using GraphPad Prism 9, which was used to generate kinetic curves and determine the Michaelis constant ($K_m$).

### Supplementary Figures

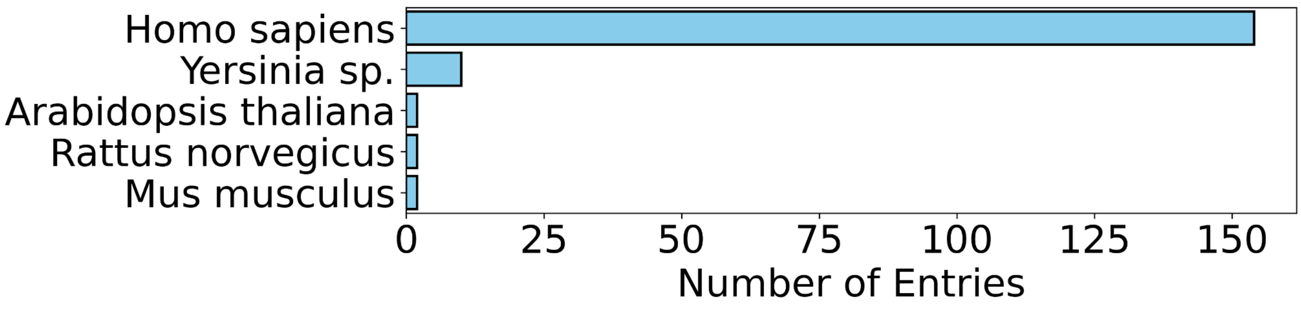


**Figure S1: Source organism distribution for PTPs.**
Bar chart showing source organisms for all PTPs used in network analysis, including mammalian, plant, and bacterial.

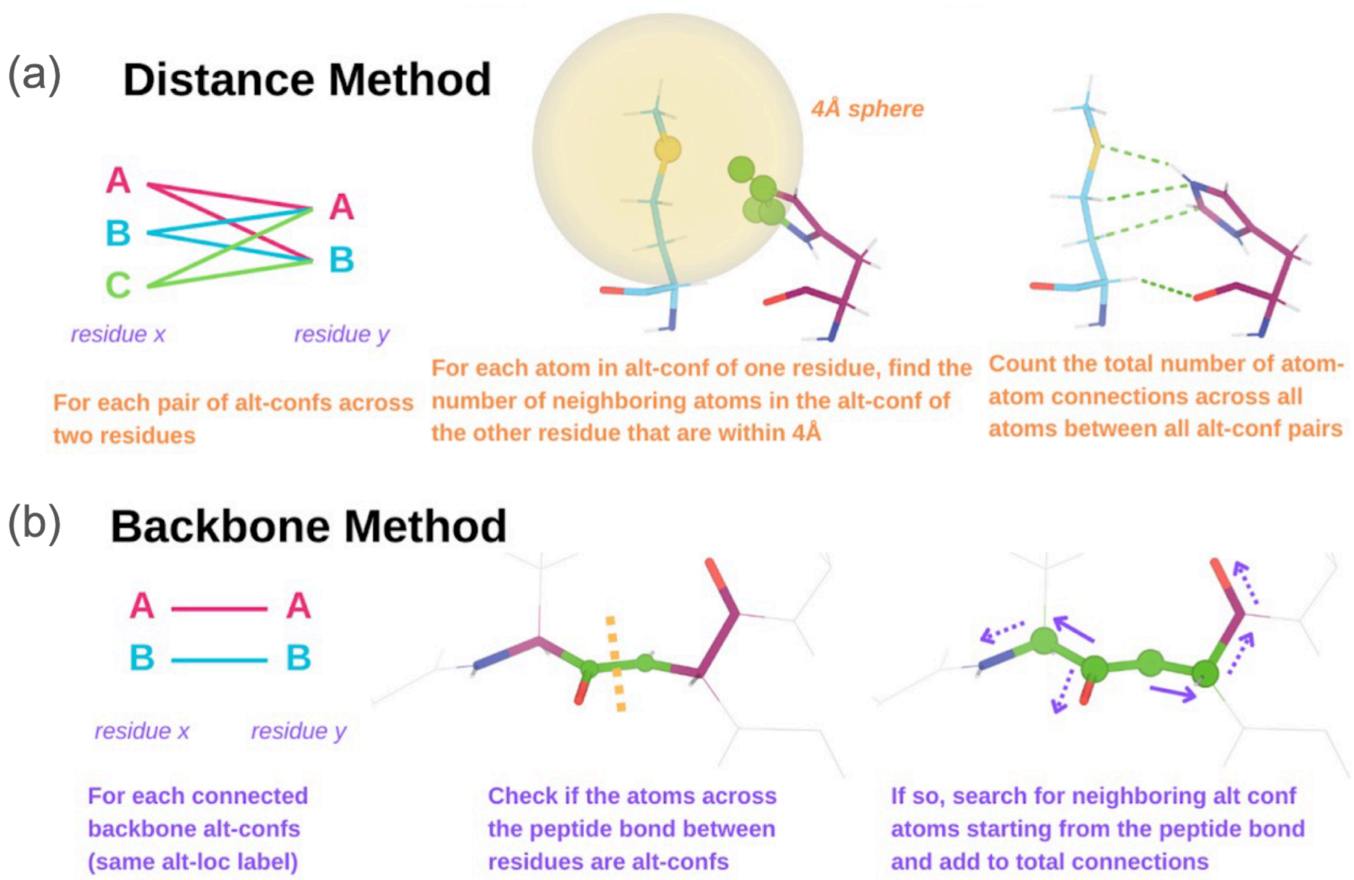


**Figure S2: Two complementary methods for defining residue-residue connections.**
The methodology used for assigning a connection/edge between two residues with alternate conformations in a RINFAIRE network, depending on whether the atoms are nearby **(a)** in space or **(b)** via covalent bonds in the protein backbone. See main **Fig. 2**.

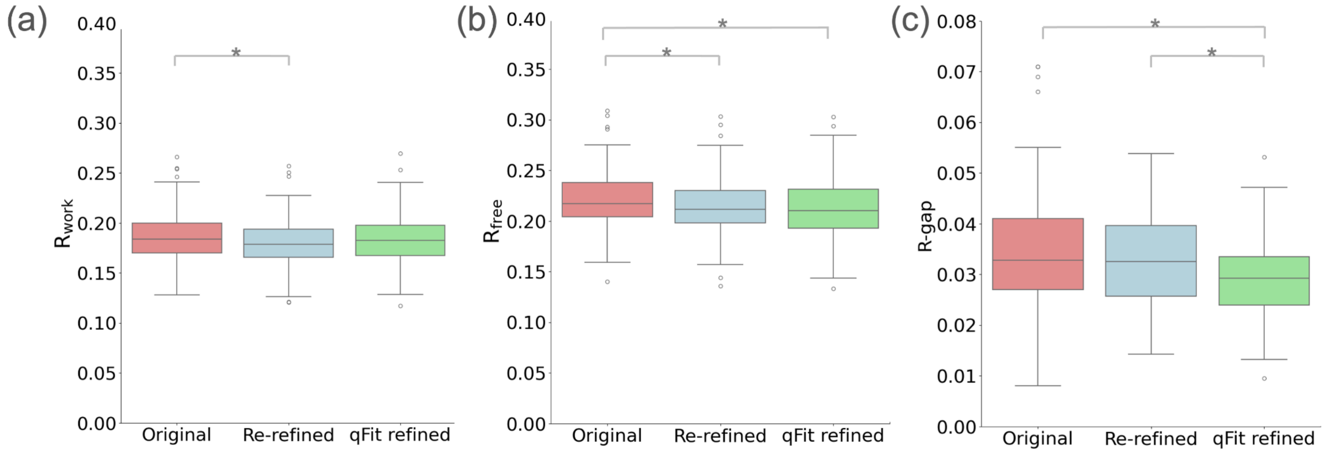


**Figure S3: Changes in R-values for automatic and qFit refinement.**
Comparison of R-factors for original, automatically re-refined, and qFit refined structures used in our analysis. Boxes represent the interquartile range (IQR), central lines represent the median, whiskers represent 1.5x the IQR, and points are outliers beyond the whiskers. * $p < 0.05$ from two-tailed Student's t-test, indicating the distributions are statistically significantly different.
**(a)** $R_{work}$ for original deposited (mean: 0.186), automatically re-refined (0.181), and qFit refined (0.183) structures.
**(b)** $R_{free}$ for original deposited (0.219), automatically re-refined (0.214), and qFit refined (0.212) structures.
**(c)** R-gap ($R_{free}$-$R_{work}$) for original deposited (0.033), automatically re-refined (0.033), and qFit refined (0.029) structures.

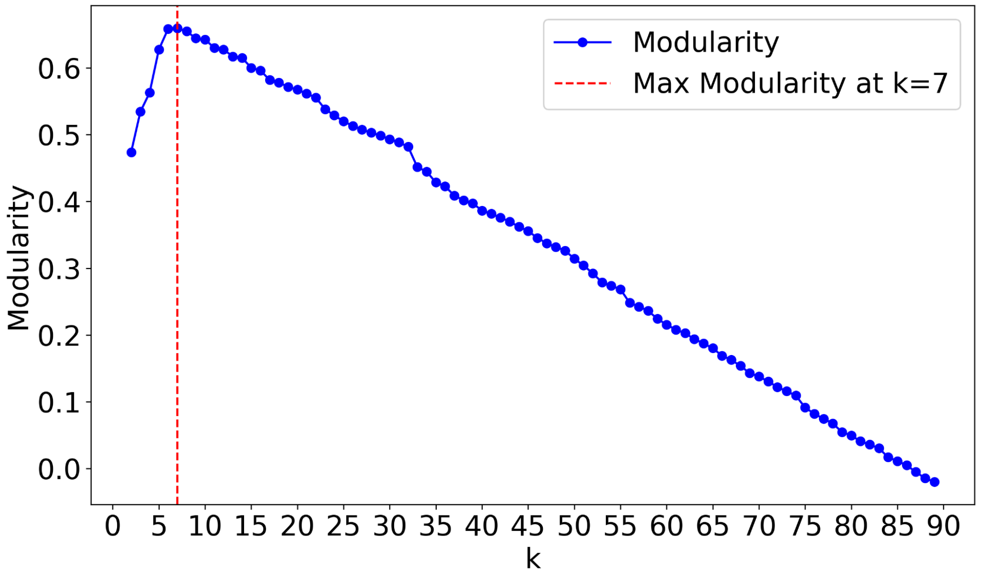


**Figure S4: Determining number of clusters based on modularity.**
Plot of modularity vs. number of clusters (k), for Girvan-Newman community detection. The red dotted line marks the value of k where modularity is maximal, indicating the optimal number of clusters. See main **Fig. 3c-d**.

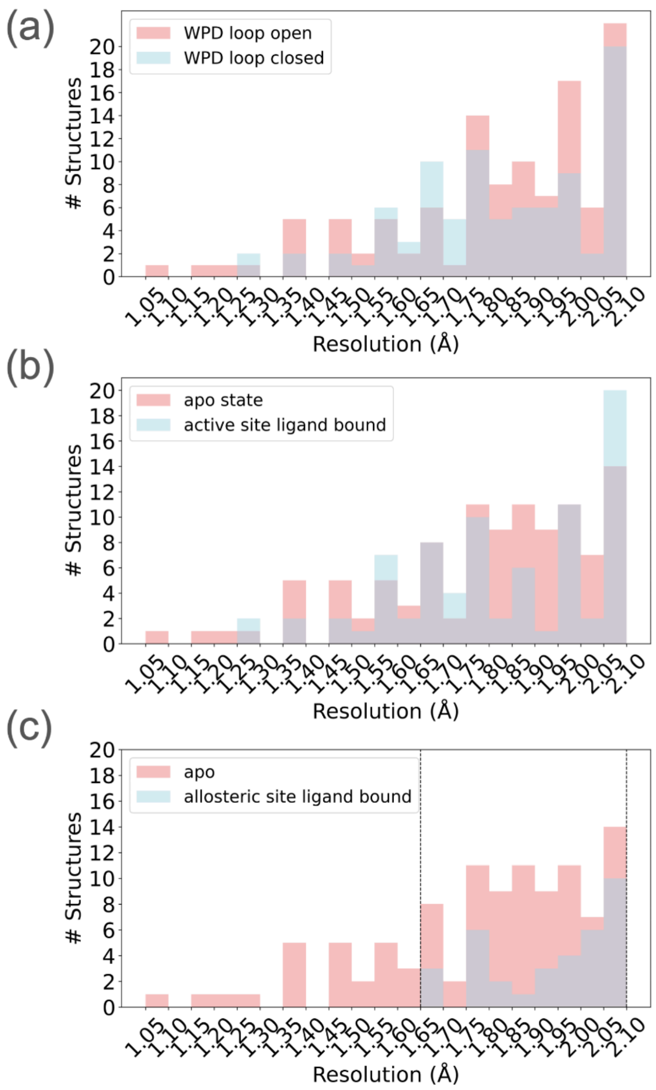


**Figure S5: Resolution distributions for WPD open/closed state and ligand-bound structures.** Resolution distributions for all PTP structures in different active-site conformations and ligand states. Histograms of resolution for each of the pairwise subsets of PTP structures used for analysis in main **Fig. 4**. For each panel, a one-tailed Mann-Whitney U test was performed to compare the two distributions. The p-value was calculated for each pair of distributions ($p < 0.05$ indicates significantly different). The dotted line (if shown) indicates that only structures within the defined resolutions were used.

**(a)** WPD loop closed vs. open. Used 1.05–2.10 Å (inclusive) resolution range; $p = 0.72$.

**(b)** Bound to active-site ligand vs. apo. Used 1.05–2.10 Å (inclusive) resolution range; $p = 0.22$.

**(c)** Bound to allosteric ligand vs. apo. Used 1.65–2.10 Å (inclusive) resolution range; $p = 0.07$.

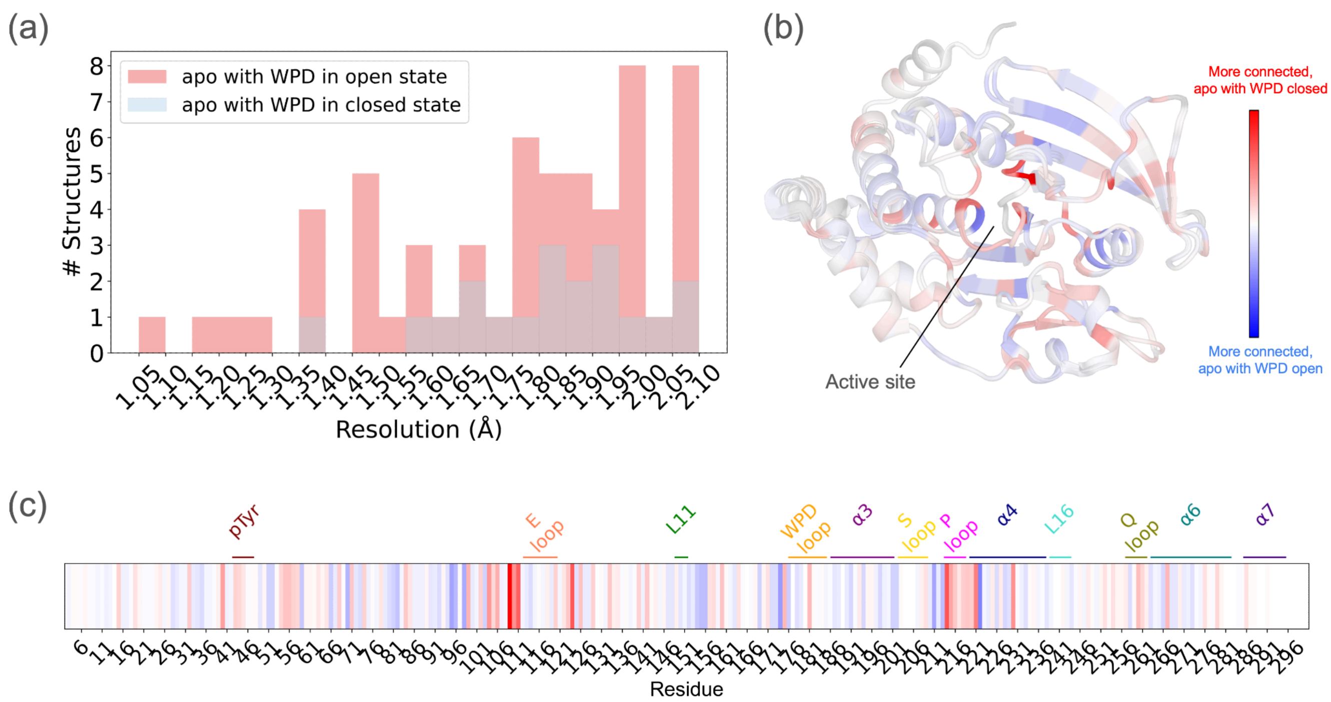


**Figure S6: Resolution distribution and sum network comparison for apo structures in WPD closed vs. open states.**

Resolution distributions and degree differences for WPD closed vs. open conformations in the apo state. Analysis of difference in weighted degree (Δdegree) for each residue in the sum networks for PTP structures in the WPD open state with no ligands vs. those with the WPD closed state with no ligands.

**(a)** Histogram of resolution for the relevant subset of structures. A one-tailed Mann-Whitney U test was performed to compare the two distributions. The p-value was calculated for the pair of distributions ($p < 0.05$ indicates significantly different). The resulting p-value was 0.51.

**(b)** Δdegree is mapped onto a cartoon visualization of structurally aligned, representative closed vs. open-state structure of the PTP catalytic domain (PDB ID: 1sug, 1t49) [18,54]. See color bar labels for red/blue coloring conventions.

**(c)** Δdegree is mapped onto a 1-dimensional representation of the protein sequence (PTP1B numbering), with key regions labeled.

Compare to main **Fig. 4a,d**.

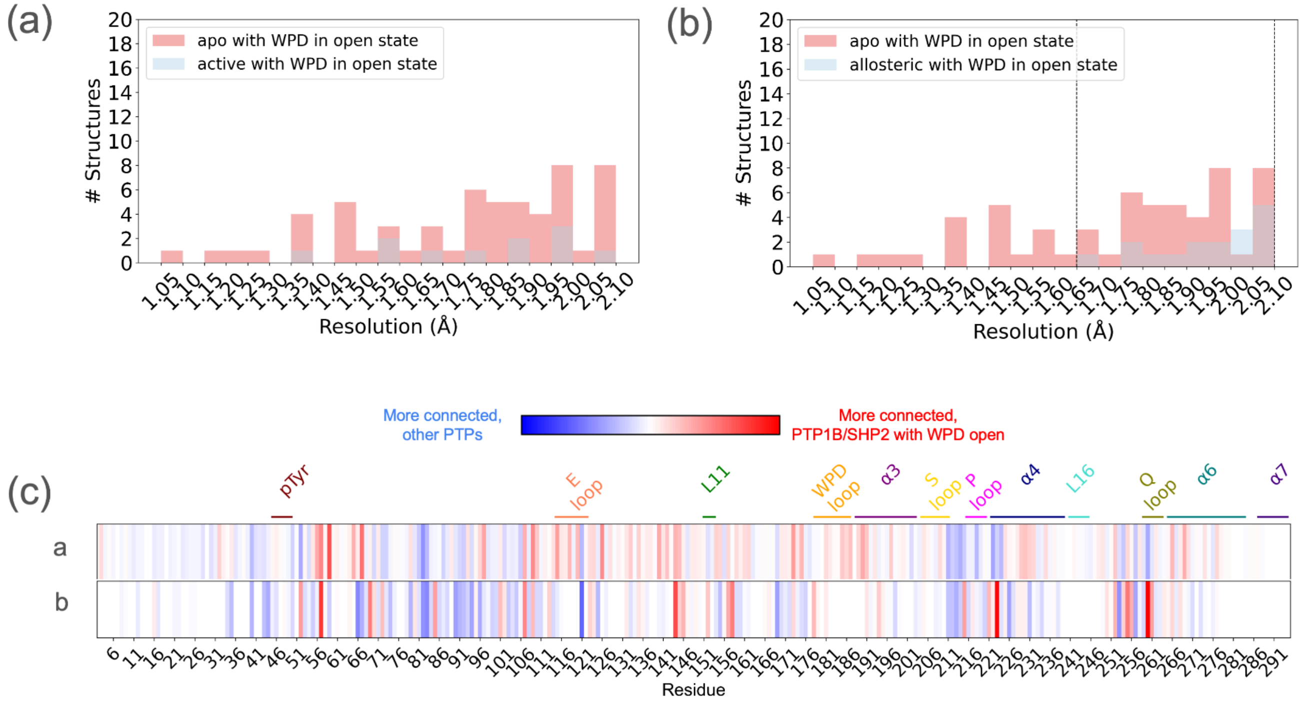


**Figure S7: Resolution distributions and sum network comparisons for ligand-bound structures in WPD open state.**

Resolution distributions and degree differences for active-site or allosteric ligands bound to a WPD open conformation. Analysis of difference in weighted degree (Δdegree) for each residue in the sum networks for the following subsets of PTP structures: active-site ligand structures in the WPD open state vs. apo structures in the WPD open state, and allosteric ligand structures in the WPD open state vs. apo structures in the WPD open state.

**(a-b)** Histograms of resolution for the relevant subsets of structures. For each panel, a one-tailed Mann-Whitney U test was performed to compare the two distributions. The p-value was calculated for each pair of distributions ($p < 0.05$ indicates significantly different). The resulting p-values were (a) 0.73 and (b) 0.13. The dotted line (if shown) indicates that only structures within the defined resolutions were used. In (b), used 1.65–2.10 Å (inclusive) resolution range.

**(c)** Δdegree is mapped onto a 1-dimensional representation of the protein sequence (PTP1B numbering), with key regions labeled. a/b labels on the left correspond to the panels in the row above.

Compare to main **Fig. 4b-d**.

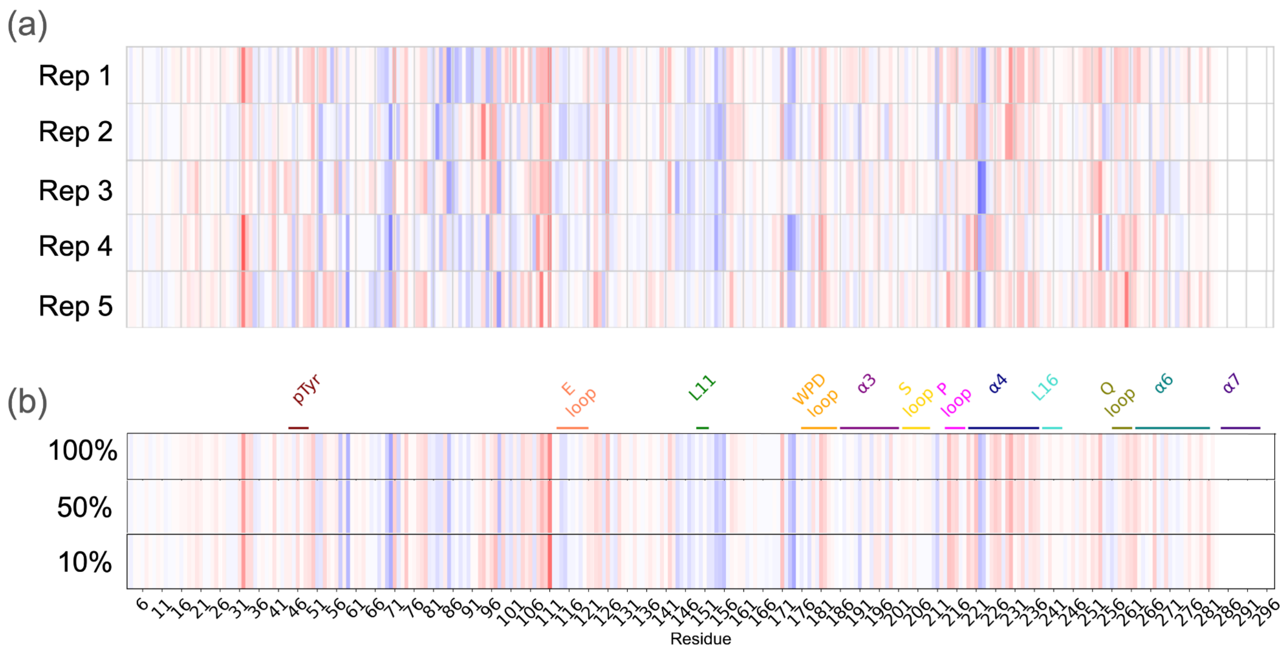


**Figure S8: Robustness of Δdegree plots to random subsetting of input structures.**
A comparison of Δdegree plots using sum networks derived from random subsets of the WPD open state vs. WPD closed state structures.
**(a)** Δdegree plots with all edges for 5 different randomly selected non-overlapping halves using 50% of all structures.
**(b)** Δdegree plots with all edges, averaged across a series of 50 random subsets using from 100% of structures to 10% of structures.

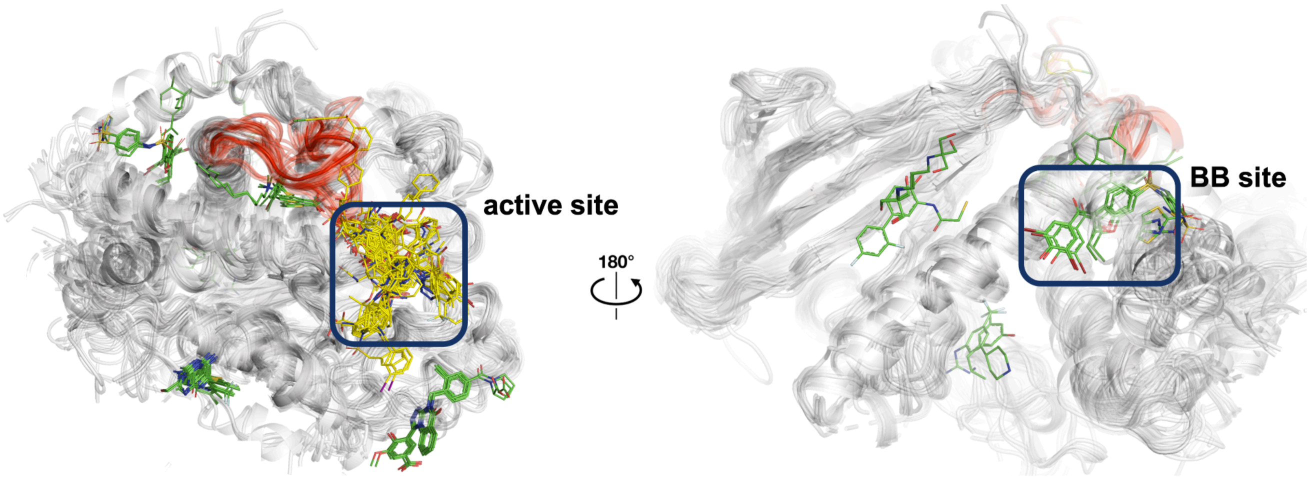


**Figure S9: All PTP structures used in the analysis with bound active-site or allosteric ligands.**
Overlay of all active-site and allosteric ligand-bound structures (individual chains aligned) used in this study. The protein is shown in gray cartoon representation. The active site and one allosteric site (BB site) are enclosed in boxes, and the WPD loop is shown in red. Active-site ligands are shown in yellow; allosteric (non-orthosteric) ligands are shown in green.

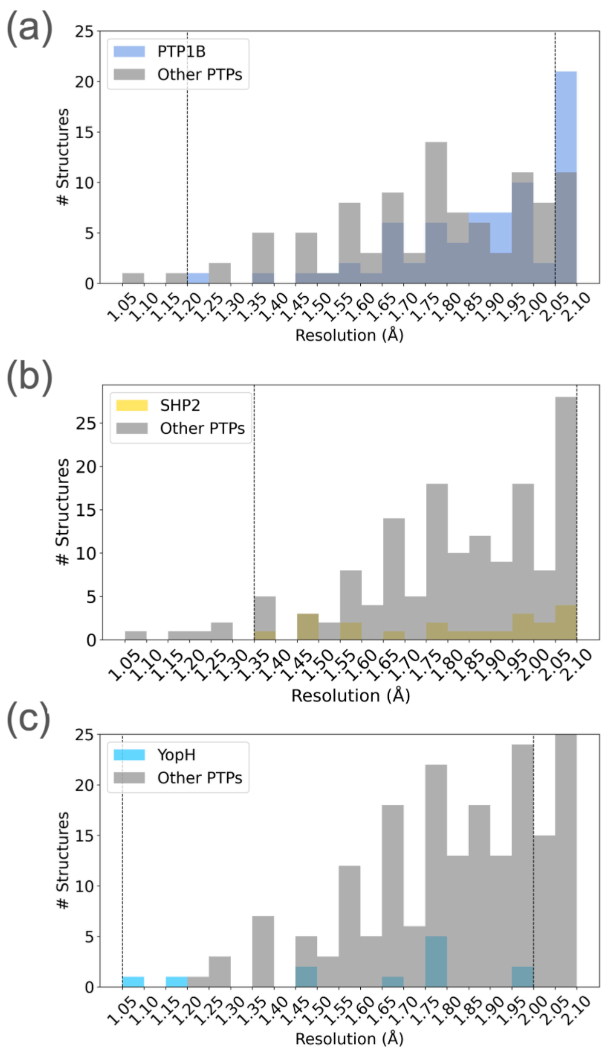


**Figure S10: Resolution distributions for PTP1B, SHP2, and YopH structures.**
Histograms of resolution for each of the pairwise subsets of PTP structures used for analysis in main **Fig. 5**. For each panel a one-tailed Mann-Whitney U test was performed to compare the two distributions. The p-value was calculated for each pair of distributions ($p < 0.05$ indicates significantly different). The dotted line (if shown) indicates that only structures within the defined resolutions were used.
**(a)** PTP1B vs. all other PTPs. Used 1.20–2.05 Å (inclusive) resolution range; $p = 0.14$.
**(b)** SHP2 vs. all other PTPs. Used 1.35–2.10 Å (inclusive) resolution range; $p = 0.76$.
**(c)** YopH vs. all other PTPs. Used 1.05–2.00 Å (inclusive) resolution range; $p = 0.16$.

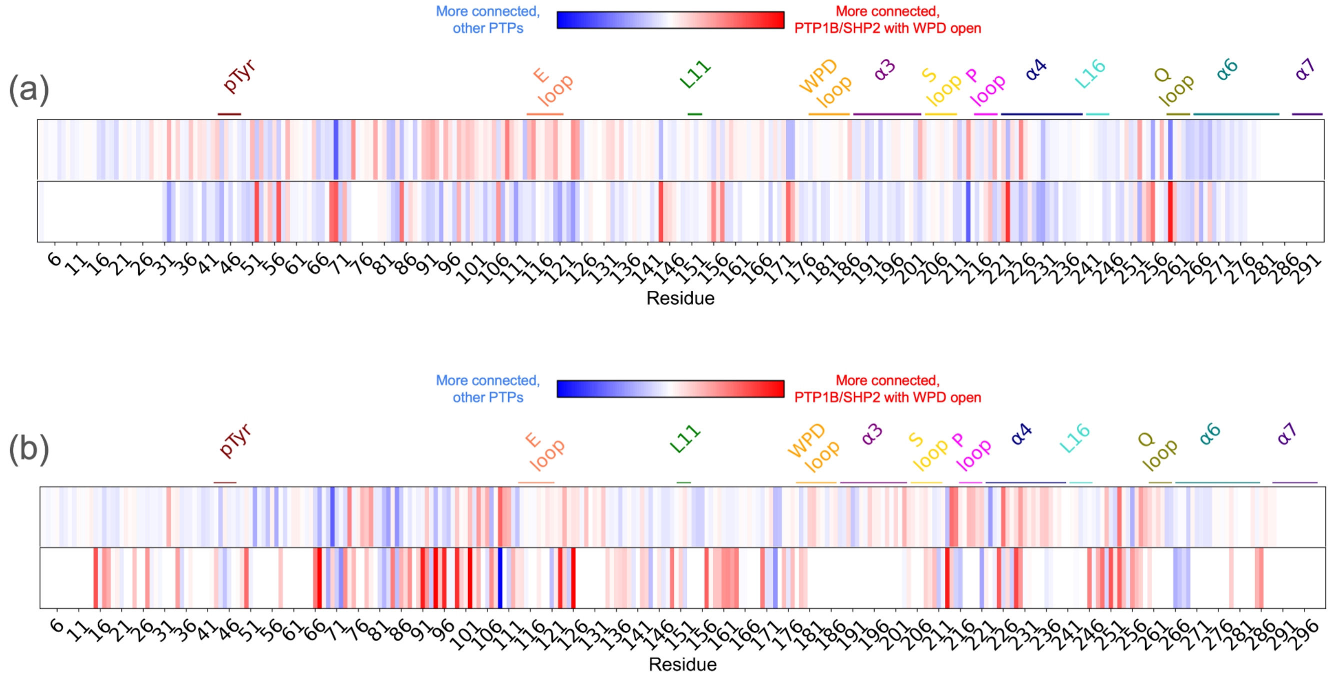


**Figure S11: Sum network comparison between PTP1B, SHP2, and YopH in the WPD open/closed state.** Analysis of difference in weighted degree (Δdegree) for each residue in the sum networks for the following subsets of PTP structures:

**(a)** PTP1B in the WPD open state vs. all non-PTP1B structures; SHP2 in the WPD open state vs. all non-SHP2 structures,

**(b)** PTP1B in the WPD closed state vs. all non-PTP1B structures; YopH in the closed state vs. all non-YopH structures.

In each case, Δdegree was calculated relative to all other available PTP structures as a reference. Δdegree is mapped onto a 1-dimensional representation of the protein sequence (PTP1B numbering), with key regions labeled.

Compare to main **Fig. 5**.

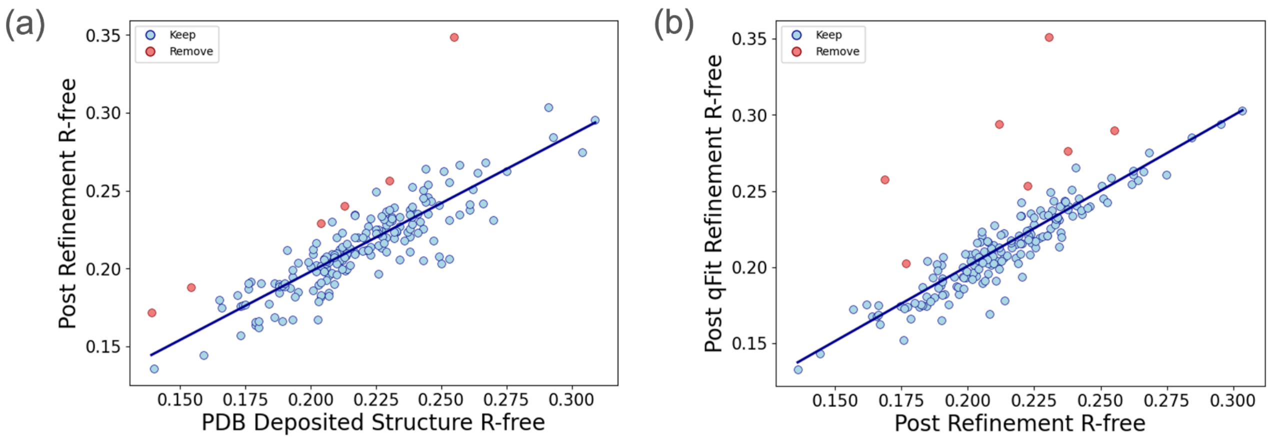


**Figure S12: $R_{free}$ values after automated re-refinement vs. qFit modeling and refinement.**
Blue dots indicate structures with a change in $R_{free} \leq 2.5\%$.
Red dots indicate structures with an increase in $R_{free} \geq 2.5\%$.
Diagonal line indicates structures with negligible change in their $R_{free}$ before and after refinement.
**(a)** $R_{free}$ values upon initial refinement vs. in the original PDB deposition.
**(b)** $R_{free}$ values post qFit refinement vs. upon initial refinement.

#### Supplementary References


106. Mistry, J. *et al.* Pfam: The protein families database in 2021. *Nucleic Acids Res.* **49**, D412–D419 (2021).

107. Headd, J. J. *et al.* Use of knowledge-based restraints in phenix.refine to improve macromolecular refinement at low resolution. *Acta Crystallogr. D Biol. Crystallogr.* **68**, 381–390 (2012).

108. Afonine, P. V., Grosse-Kunstleve, R. W., Adams, P. D. & Urzhumtsev, A. Bulk-solvent and overall scaling revisited: faster calculations, improved results. *Acta Crystallogr. D Biol. Crystallogr.* **69**, 625–634 (2013).

109. Afonine, P. V. *et al.* Towards automated crystallographic structure refinement with phenix.refine. *Acta Crystallogr. D Biol. Crystallogr.* **68**, 352–367 (2012).

110. Pei, J., Tang, M. & Grishin, N. V. PROMALS3D web server for accurate multiple protein sequence and structure alignments. *Nucleic Acids Res.* **36**, W30–4 (2008).

111. Hagberg, A. A., Schult, D. A. & Swart, P. J. Exploring network structure, dynamics, and function using NetworkX. in *Proceedings of the Python in Science Conference* 11–15 (SciPy, 2008).

112. Hongdusit, A., Zwart, P. H., Sankaran, B. & Fox, J. M. Minimally disruptive optical control of protein tyrosine phosphatase 1B. *Nat. Commun.* **11**, 788 (2020).

113. Shen, R. *et al.* Insights into the importance of WPD-loop sequence for activity and structure in protein tyrosine phosphatases. *ChemRxiv* (2022) doi:10.26434/chemrxiv-2021-8f6mc-v2.

114. Chu, H.-M. & Wang, A. H.-J. Enzyme-substrate interactions revealed by the crystal structures of the archaeal Sulfolobus PTP-fold phosphatase and its phosphopeptide complexes. *Proteins* **66**, 996–1003 (2007).

115. Wojdyr, M. GEMMI: A library for structural biology. *J. Open Source Softw.* **7**, 4200 (2022).

116. Valdar, W. S. J. Scoring residue conservation. *Proteins* **48**, 227–241 (2002).

117. Schrödinger, L. & DeLano, W. PyMOL. (2020).

118. Sievers, F. & Higgins, D. G. Clustal Omega for making accurate alignments of many protein sequences. *Protein Sci.* **27**, 135–145 (2018).

119. Cui, D. S., Lipchock, J. M., Brookner, D. & Loria, J. P. Uncovering the molecular interactions in the catalytic loop that modulate the conformational dynamics in protein tyrosine phosphatase 1B. *J. Am. Chem. Soc.* **141**, 12634–12647 (2019).

120. Greisman, J. B. *et al.* Native SAD phasing at room temperature. *Acta Crystallogr. D Struct. Biol.* **78**, 986–996 (2022).
